# metaGEM: reconstruction of genome scale metabolic models directly from metagenomes

**DOI:** 10.1101/2020.12.31.424982

**Authors:** Francisco Zorrilla, Kiran R. Patil, Aleksej Zelezniak

**Affiliations:** Division of Systems and Synthetic Biology, Department of Biology and Biological Engineering, Chalmers University of Technology, Gothenburg, Sweden; Structural and Computational Biology Unit, European Molecular Biology Laboratory, Heidelberg, Germany; Medical Research Council Toxicology Unit, University of Cambridge, Cambridge, UK

## Abstract

Advances in genome-resolved metagenomic analysis of complex microbial communities have revealed a large degree of interspecies and intraspecies genetic diversity through the reconstruction of metagenome assembled genomes (MAGs). Yet, metabolic modeling efforts still tend to rely on reference genomes as the starting point for reconstruction and simulation of genome scale metabolic models (GEMs), neglecting the immense intra- and inter-species diversity present in microbial communities. Here we present metaGEM (https://github.com/franciscozorrilla/metaGEM), an end-to-end highly scalable pipeline enabling metabolic modeling of multi-species communities directly from metagenomic samples. The pipeline automates all steps from the extraction of context-specific prokaryotic GEMs from metagenome assembled genomes to community level flux balance simulations. To demonstrate the capabilities of the metaGEM pipeline, we analyzed 483 samples spanning lab culture, human gut, plant associated, soil, and ocean metagenomes, to reconstruct over 14 000 prokaryotic GEMs. We show that GEMs reconstructed from metagenomes have fully represented metabolism comparable to the GEMs reconstructed from reference genomes. We further demonstrate that metagenomic GEMs capture intraspecies metabolic diversity by identifying the differences between pathogenicity levels of type 2 diabetes at the level of gut bacterial metabolic exchanges. Overall, our pipeline enables simulation-ready metabolic model reconstruction directly from individual metagenomes, provides a resource of all reconstructed metabolic models, and showcases community-level modeling of microbiomes associated with disease conditions allowing generation of mechanistic hypotheses.

## Introduction

Whole metagenome shotgun sequencing and genome-resolved metagenomics allow for the exploration of personalized, context-specific microbial communities at a species or strain level resolution^1^. Changes of microbiota composition are strongly linked to a range of diseases including cancer, behavioral, neurological, and metabolic disorders^2–7^. However, a mechanistic understanding of the role of the human gut microbiome in disease, especially the roles of specific strains and the associated metabolic factors, remains challenging due to the vast intra- and inter-species diversity. To this end, short-read sequencing data allows for the extraction of metagenome assembled genomes (MAGs) directly from raw sequencing data, while avoiding culture-based methods or making use of the limited number of reference genomes, enabling the discovery of unknown species and the exploration of personalized microbiomes^8–10^. Indeed, a number of attempts aiming to explore the human gut microbiome composition and diversity have generated hundreds of MAGs representing previously unknown or uncultured species, as well as thousands of known species MAGs^11–13^. Pangenome analysis of the human gut microbiome demonstrated that the functional repertoire of gut species differ significantly, with a median core genome proportion of only 66%^14^, revealing differences in metabolic potentials of individual microbiomes.

Attempts of mechanistic links between diet or inter-species interactions with microbiota composition and dynamics lead to development of gut species genome-scale metabolic models^15^. GEnome-scale Metabolic models (GEMs) allow assessing species nutritional requirements^16^ and their interactions in the human gut as well as in diverse environmental communities^17–19^. The current paradigm of metabolic modeling typically relies on mapping identified taxa to their closest reference genomes. This limits analysis and its interpretation to the metabolic networks represented in the known reference genome space. This can cause false positives (i.e. pathways present in the reference but missing from the variant present in the community) as well as false negatives (i.e. pathways missing in the reference genomes but present in the community variant), ultimately leading to inaccurate predictions of individual species metabolism as well as that of cross-feeding interactions. Thus the current modelling attempts are likely failing to capture the specific metabolic features of a given species across different contexts, e.g. microbiota of individuals with different disease conditions. Towards overcoming this limitation, here we present metaGEM, the computational pipeline that enables reconstruction of sample specific metabolic models directly from short read metagenomics data. Instead of relying on reference genomes, metaGEM generates high quality metagenome assembled genomes, which are then used to reconstruct context-specific prokaryotic GEMs using state-of-the-art methodologies (Figure 1). Our contributions are two-fold: i) an end-to-end framework enabling community-level metabolic interaction simulations directly from metagenomes, and ii) a resource of more than 14 000 MAGs from a range of metagenomic biomes, including 3750 high quality MAGs, with corresponding ready-to-use GEMs from human gut microbiome studies^20,21^ and global microbiome projects^22–24^. The metaGEM pipeline is implemented in Snakemake^25^, an open-source, community driven, and scalable bioinformatics workflow engine, supporting major popular high-performance-computing cluster environments as well as standalone systems (Supplementary figure 1).

**Fig. 1:**
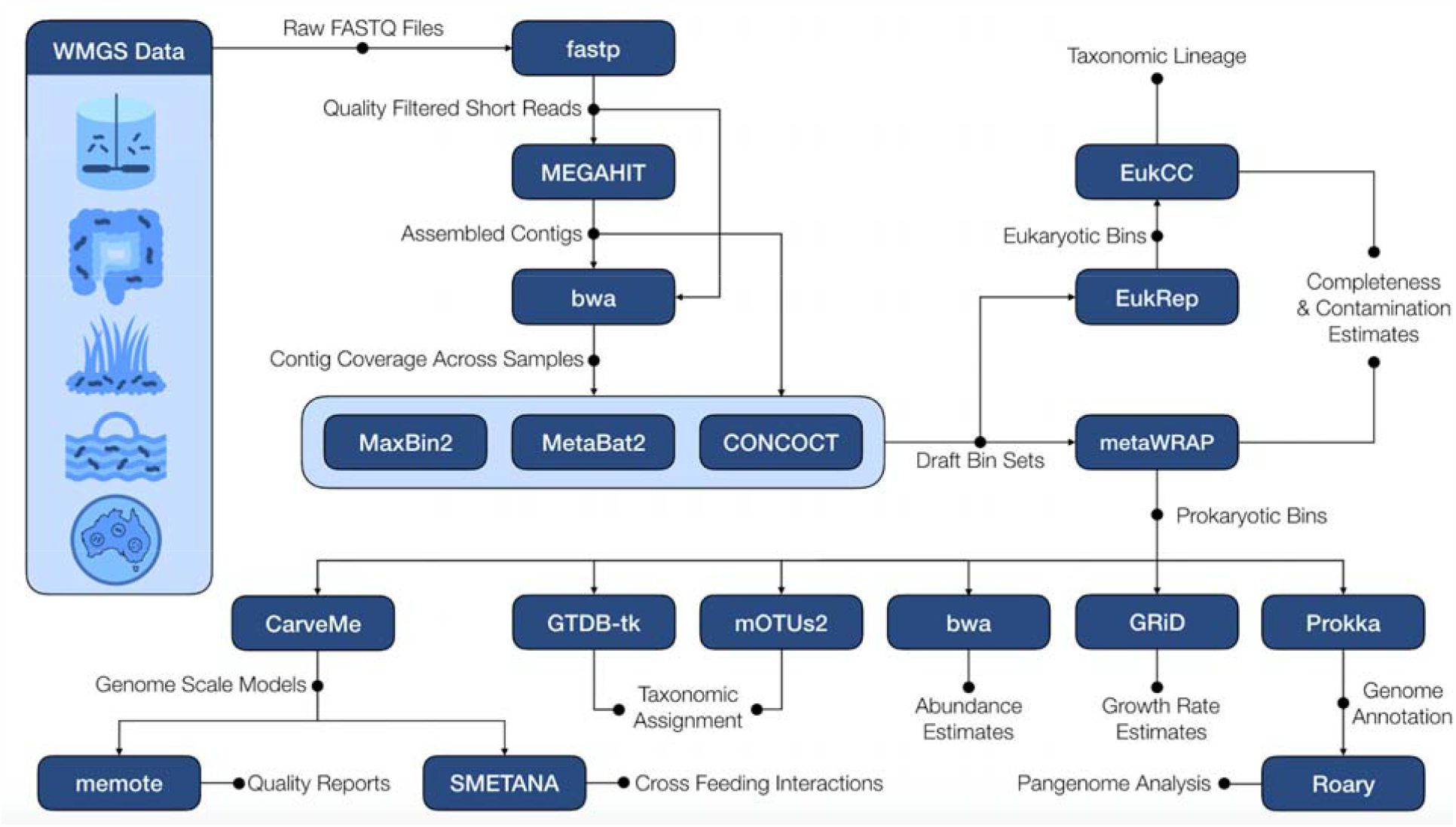
Schematic of the metaGEM pipeline workflow highlighting tools, inputs, and outputs. Short reads are quality filtered and adapter trimmed using fastp^43^. Quality controlled reads are assembled individually using MEGAHIT^44^. Using either kallisto^45^ or bwa^46^, quality controlled reads are mapped to each assembly to obtain contig coverage information across samples. Coverage information and assemblies are used by CONCOCT^8^, MetaBat2^10^, and MaxBin2^9^ to generate three bin sets for each sample. The metaWRAP^26^ bin_refinement module is used to consolidate dereplicate bin sets for each sample and find the highest quality version of each bin. The metaWRAP bin_reassemble module is used to extract quality controlled short reads mapping from the focal sample to the bin, which are used to generate two single genome assemblies using strict and permissive parameters. The original and reassembled versions are compared for quality, and the best verison is kept. Refined and reassembled MAGs are used to generate GEMs using CarveMe^28^. These models can be quality checked using memote^29^. Community simulations are carried out for each sample using SMETANA^17^. Other features include: taxonomic classification using mOTUs2^47^ and/or GTDB-Tk^48^, a custom mapping-based abundance estimation module that does not make use of marker-gene or reference-genome based approaches, growth rate estimation for high coverage MAGs using GRiD^30^, and pangenome analysis using prokka^49^ and roary^31^. EukRep^50^ can be used to scan for eukaryotic contigs in the CONCOCT bin sets, which can then be processed by EukCC^51^ to provide completeness, contamination, and taxonomic assignments for eukaryotic MAGs.

## Results

### Implementation and features of metaGEM pipeline

The metaGEM pipeline starts from the single sample assembly of short read FASTQ data, and proceeds to reconstruction of metagenome assembled genomes using three different binners: CONCOCT^8^, MetaBat2^10^, MaxBin2^9^. The three output draft bin sets are then refined (i.e. de-replicated) and reassembled using metaWRAP to obtain the highest quality version of each bin^26^ (Supplementary figure 2). Note that metaWRAP completeness and contamination estimates are based on a marker-gene approach used by CheckM^27^. These final prokaryotic MAGs are used to automatically generate flux balance analysis ready genome-scale metabolic reconstructions using CarveMe^28^, which are quality checked using the memote^29^ framework. By integrating the Species METabolic ANAlysis (SMETANA) framework^17^ (https://github.com/cdanielmachado/smetana), the generated GEMs can be used for sample specific community-level metabolic interaction modeling. Other pipeline features include the automatic assignment of taxonomic classification to the reconstructed MAGs and GEMs, the calculation of relative and absolute abundance for generated MAGs, the estimation of growth rate^30^ for high coverage MAGs, and pangenome analysis^31^ (Figure 1). Although there is a growing number of MAG reconstruction pipelines^32–38^, a simple comparison of studies that used differing methods for MAG generation revealed that metaGEM consistently recovers more high quality genomes per sample, both from gut microbiome^11–13^ and ocean^39–41^ metagenomes (Supplementary figure 3). Indeed, metaGEM has already been applied in one of our recent studies to interrogate the plastic-degrading potential of the global microbiome^42^.

To demonstrate the versatility of metaGEM pipeline, we reconstructed genome-scale metabolic models (GEMs) from five metagenome datasets spanning different biological and technical complexity, namely: gut microbiomes^20,21^, plant associated microbiomes^22^, global ocean microbiomes^23^, and bulk soil microbiomes^24^. In total we reconstructed 3750 high quality metagenomic assemblies (MAGs) with >90% completeness & <5% contamination and 10349 medium quality MAGs with >50% completeness and <5% contamination (Supplementary figure 4). We assessed the quality of MAGs reconstruction by recovering genomes from a controlled multispecies lab culture experiment^20^. Briefly, 7 human gut microbiome species were grown in vitro across 4 biological replicates with 12 time points totaling 48 metagenomic samples. A total of 154 MQ MAGs, of which 137 also meet the HQ MAG standard, were reconstructed with an average completeness of 95.4% and average contamination of 0.3%, with on average ∼3.2 MAGs per sample reflecting the growth curves shown in the original publication are dominated by very few species early on and ultimately just one (Supplementary figure 5). Abundance estimates generated using a mapping-based approach (Methods) were perfectly correlated to marker gene based abundance estimates (Pearson’s *r* = 0.99, p-value <1e-16) well recapitulating experimental observations (Figure 2a).

**Fig. 2:**
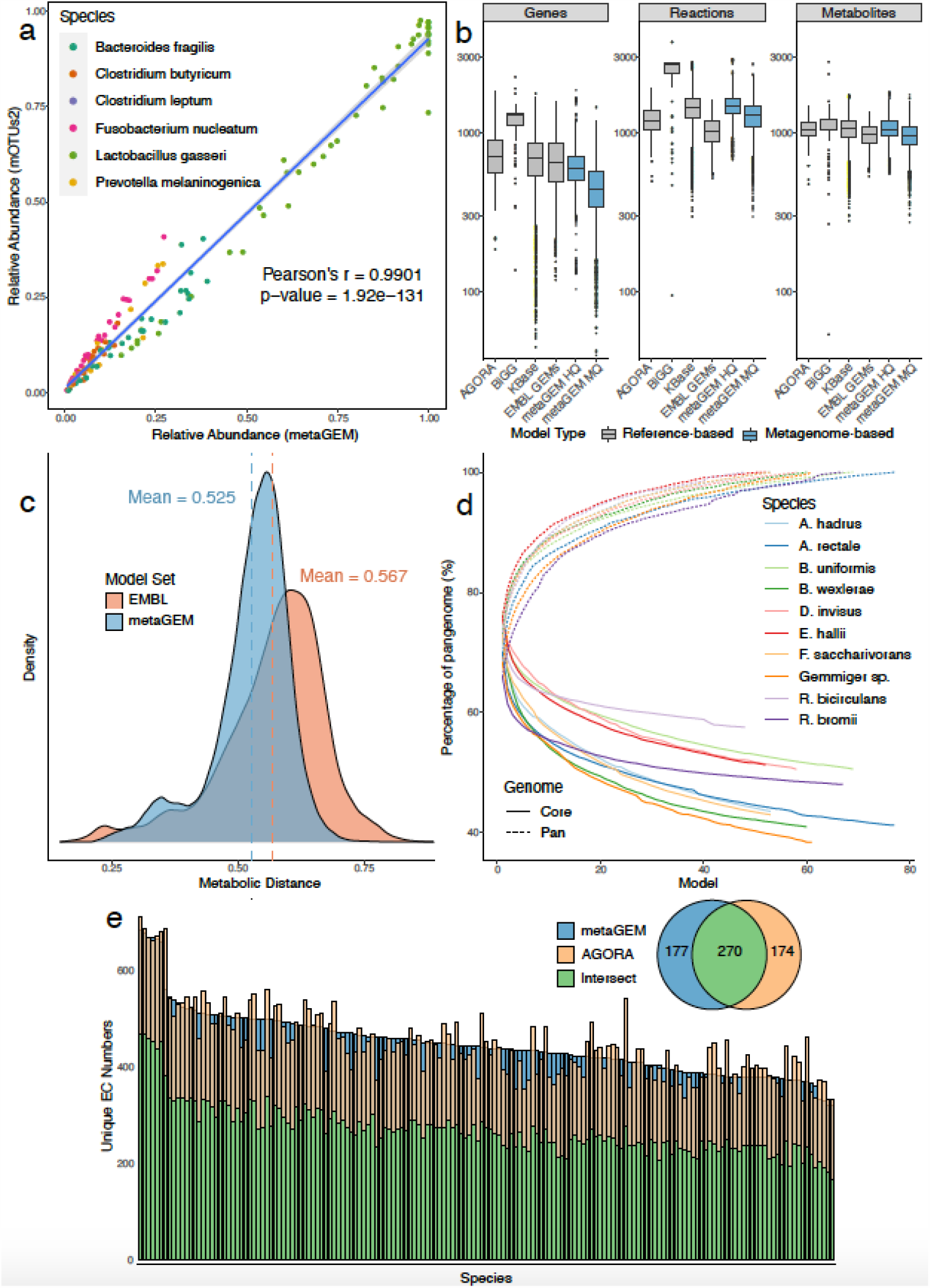
Abundance, quality, and diversity comparisons of reconstructions. (a) Abundance estimates generated by metaGEM using a mapping based approach compared to marker gene based approach of mOTUs2 in small lab culture communities dataset. (b) Distribution of genes, reactions, and metabolites in genome scale metabolic models across AGORA^15^, BiGG^52^, EMBL GEMs^28^, KBase^38^, medium quality (MQ) metaGEM, and high quality (HQ) metaGEM sets. (c) Distribution of metabolic distances between a set of 200 randomly chosen reference EMBL GEMs compared to a randomly chosen set of 800 EMBL GEMs and 800 randomly chosen metaGEMs from the gut microbiome dataset. (d) Cumulative core and pan genome curves for the top ten most commonly reconstructed gut microbiome species based on EC numbers present in the reconstructed genome scale metabolic models. (e) Comparison of EC numbers between 165 species reconstructed from the gut microbiome dataset and also found in the AGORA collection. Inset venn diagram shows average value of EC numbers unique to the compared sets as well as their average intersect.

### High-quality metabolic reconstructions directly from metagenomes

We next reconstructed a total of 14087 GEMs from the metagenome assembled genomes (MAGs) (Methods) and compared them to highly-curated reference genome-based BiGG models^52^, AGORA^15^, EMBL^28^ and KBase models^38^. In terms of the number of metabolic reactions and unique metabolites, the metagenomic GEMs show similar distributions compared to reference-based GEM reconstructions (Figure 2b). Specifically, high quality metagenome derived GEMs were on average 101.6%, 99.7%, and 110.0% similar in terms of the number of metabolites and 103.7%, 121.1%, 144.3% similar in terms of the number of reactions respectively compared to reference genome-based EMBL, AGORA and KBase models, with 95.9% less metabolites and 59.9% less reactions compared to manually curated BiGG models, which overall display a higher number of reactions and metabolites. In terms of the number of genes present in the models, the high quality metagenome derived GEMs had 9.5% less genes as compared to the reference genome-based collections, with the exception of BIGG models that involved over 40% more gene annotations. On the contrary, metaGEM models were on average 79.7%, 0.5%, 0.7%, and 0.5% similar in terms of the number of dead end metabolites compared to EMBL, AGORA, KBase, and BiGG models, respectively (Supplementary figure 6). Furthermore, reconstructed metaGEMs using high quality MAGs and medium quality MAGs were 75.4%, 87.2%, and 90.7% similar in terms of their number of genes, reactions, and metabolites, suggesting that the metabolic reconstruction process^28^ is robust towards bin completion. Separating metagenomic models by dataset shows that the distributions of metabolites, reactions, and genes are similar respectively across datasets as well (Supplementary figure 7).

We also evaluated whether the metabolic reaction diversity identified in metagenomes would be comparable to the expected enzymatic diversity from the reference genome reconstructions, i.e. if expected reactions and pathways would be present in metagenomes based GEMs. For this we randomly sampled 1000 metabolic models from each, genome-based and metagenome-based reconstructions, and computed metabolic similarity (expressed as Jaccard index) to the each mode of random 20% held-out set (Figure 2c). Metagenomic GEMs and reference genome based EMBL GEMs had an average had a negligible 4.2% difference (Wilcoxon rank sum test p-value < 2.2e-16) in metabolic reactions compared to an independent subset of metabolic reconstructions from reference based genomes, suggesting that metabolic models derived from metagenome assembled genomes capture expected metabolic diversity and features as reconstructions from reference-based genomes (Methods). Performing PCA on the presence/absence of EC numbers across models resulted in AGORA and gut metagenomic models clustering near each other, while EMBL and ocean metaGEM models clustered closer to each other (Supplementary figure 8).

### Pan-metabolism of metagenomic GEMs uncovers metabolic diversity within species

Recent gut metagenomic microbial population studies suggest that strain-level diversity within the same species can differ by over 20% of genome content between individuals^53^. With more extreme examples from environmental isolates of soil myxobacteria with only 30% conserved core part of genomes showing extreme gene diversity within the same species ^54^. These analyses imply that current reference based approaches^15^ may not always reflect the metabolic diversity found across a species’ pangenome or pan-metabolism. To investigate whether reconstructed metagenomic GEMs are able to capture this intra-species metabolic diversity we compared intra-species EC numbers (i.e. the pan-metabolism) diversity across the top ten most prevalent species (Figure 2c). Pan-metabolism analysis confirmed that no two models were exactly the same with respect to their EC number content. Indeed, the core genome of metabolism in the analyzed species ranged between 38.5-57.6% of their respective pangenomes, in line with previously reported degree of intraspecies genetic variation in prokaryotes^14,53^. We also analyzed the pangenome of 141 taxonomically undefined species, where we found the core genome to be 6.9% of the pan-species pan-genome (Supplementary figure 9). To exemplify this further, we compared unique EC numbers between metagenome based GEMs and reference genome based gut AGORA models^15^ for a total of 165 matched gut microbiome species. The intersect of AGORA and metaGEM EC numbers range between 48.9% to 69% for all matched species, with metaGEM models containing more EC numbers than their corresponding AGORA model in 53.9% of cases. Inspection of the KEGG pathway annotations of the EC numbers present exclusively in AGORA or metagenomic gut models reveals that the majority of these enzymes are associated with biosynthesis of secondary metabolites and antibiotics (Supplementary figure 10).

### metaGEMs enables modeling of personalized human gut communities

To investigate potential microbial metabolic interactions in healthy and metabolically impaired type 2 diabetes human gut microbiomes^21^, we reconstructed a total of 4127 of personalized human gut metaGEMs across 137 metagenomes that were classified according to the disease condition of participants from the original study, i.e. normal glucose tolerance (NGT, n = 42), impaired glucose tolerance (IGT, n = 42), or type 2 diabetic (T2D, n= 53). We then applied Species Metabolic Coupling Analysis (SMETANA), a constrained based technique for modeling interspecies dependencies in microbial communities^17^, to elucidate the potential microbial metabolic interactions in healthy and metabolically impared patients. Briefly, SMETANA outputs a table of scores for each community, corresponding to measures of strength of cross-feeding interactions that should take place to support the growth of community members in a given condition, i.e. a likelihood of species A growth depending on metabolite X from species B.

Specifically, we found 22 compound exchanges that were significantly different (Wilcoxon rank sum test, BH adjusted p-values < 0.01) between the disease groups, representing metabolites from multiple classes including organic acids and lipid-like molecules (Supplementary figures 11 and 12). Additionally, we identified 27 donors and 27 receivers that had statistically significant distributions of SMETANA scores between disease groups (Wilcoxon rank sum test, BH adjusted p-value < 0.0001), including genera that have been associated with T2D^55^ (Supplementary figures 13 and 14) such as *Bifidobacterium, Faecalibacterium, Roseburia, Ruminococcus*, and *Blautia*. Inter-species metabolic exchanges were on average 102.9% and 103.4% different in T2D and IGT patients respectively, expressed as coefficient of variation (CV = σ/μ) of SMETANA scores. Visualization of ten compounds with the lowest p-values exchanged by the eight most frequent donors or receivers of these metabolites shows notable differences in metabolic architecture across conditions (Figure 3a). For example, in the visualized data subset, exchanges of L-Malic acid undecaprenyl diphosphate were observed 2.25 and 2.75 times more frequently, and with a 115-fold and 3.9-fold higher average SMETANA score (Wilcoxon rank sum test, BH adjusted p-value = 1.8e-05 and 1.0e-03 respectively) respectively in T2D communities compared to NGT. Although exchanges of nitrite and hydrogen sulfide were observed 1.6 and 1.4 times more frequently in T2D compared to NGT communities, exchanges in the latter had a 5.5-fold and and 5.7-fold higher SMETANA score respectively compared to T2D communities (Wilcoxon rank sum test, BH adjusted p-value = 3.3e-05 and 5.8e-04 respectively). Visualization of average SMETANA scores grouped by broad metabolite class for interactions involving *Faecalibacterium prausnitzii C* as a receiver (Figures 3b,c) also highlight differences in metabolism, with a 3.7-fold stronger dependency on organic oxygen compounds and a 2.2-fold stronger dependency on nucleosides, nucleotides, and analogues in NGT communities compared to T2D. Also of note is the fact that *Faecalibacterium prausnitzii* did not demand any inorganic compounds in NGT communities, while overall it had higher dependency on inorganic compounds (nitrite) in IGT communities compared to T2D.

**Fig. 3:**
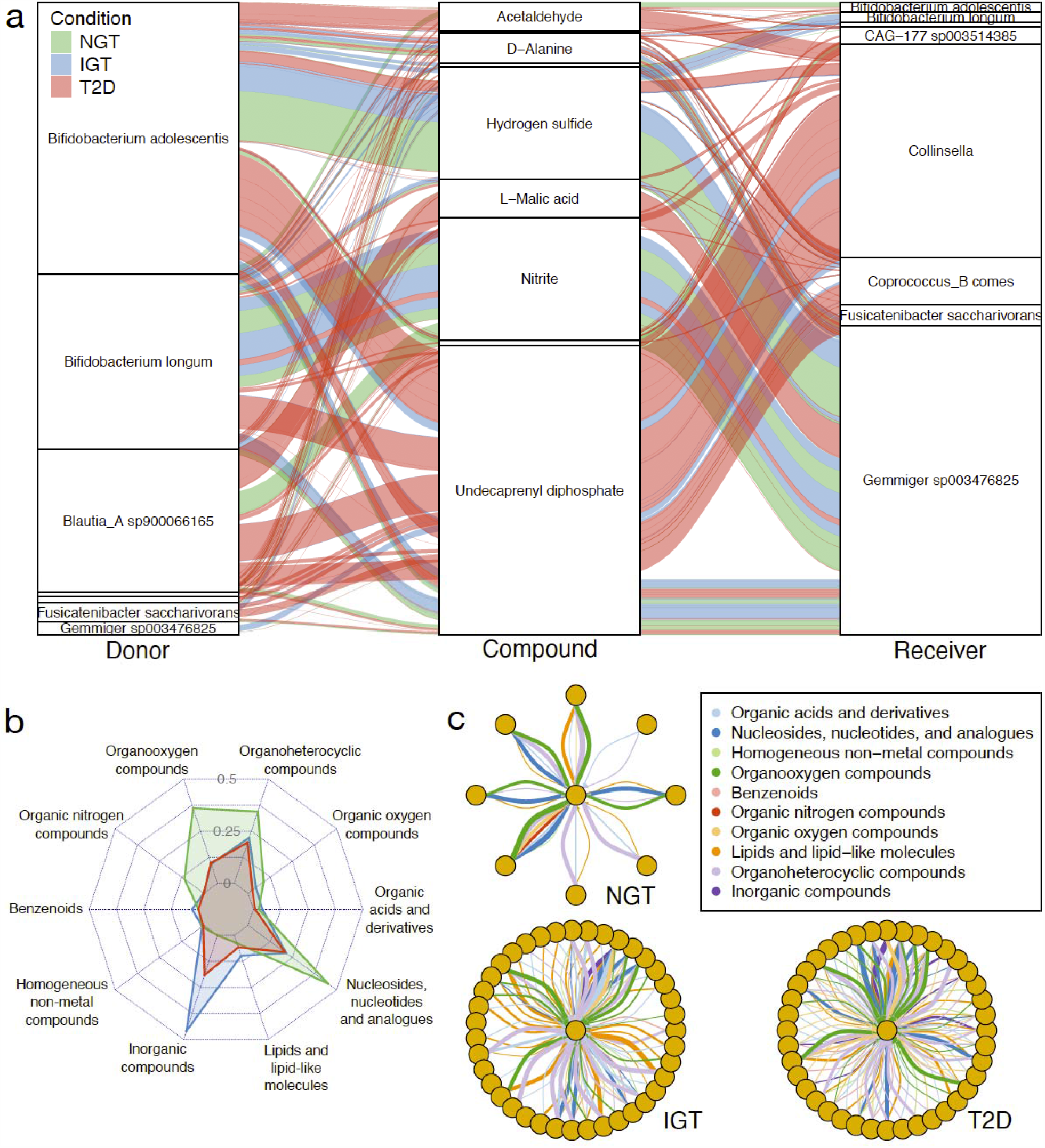
SMETANA simulations uncover differences in metabolism across conditions. a) Alluvial diagram showing top 10 compounds exchanged with statistical significance across conditions between 8 species, representing 279 interactions (NGT n=61, IGT n=58, T2D n=160) across 41 samples (NGT n=12, IGT n=12, T2D n=17). Thickness of lines are proportional to magnitude of SMETANA score. b) Radar plot of average SMETANA scores based on 543 interactions (NGT n=50, IGT n=161, T2D n=249) across 18 samples (NGT n=4, IGT n=6, T2D n=8) grouped by metabolite class across conditions for receiver *Faecalibacterium prausnitzii C*. c) Network diagrams of interactions involving *Faecalibacterium prausnitzii C* (centered in each subgraph) as a receiver across conditions. Thickness of lines are proportional to magnitude of SMETANA score.

## Discussion

The latest genome sequencing studies suggest that bacterial genomes are highly dynamic^56^, exhibiting a high exchange of genetic material between strains of the same species or between different bacterial species. With multiple examples where specific strains play key roles in disease pathogenesis^57^ or niche metabolic adaptation^54^, it becomes apparent that bacteria should be analyzed by the context specific functional repertoire (including metabolism) of strains and not merely by their species membership^58^. The highly diverse pangenome-derived functional repertoire^59,60^ of microbial communities, the interpersonal differences in gene content of human gut bacteria^53^, and the strain specific plasticity of metabolic adaptation^54^ cannot be captured by either amplicon sequencing nor by using reference based genomes. To address these limitations, and to further enable the interrogation of functional and metabolic diversity existing within microbial communities derived from metagenomes, here we developed metaGEM, a pipeline for reconstruction and metabolic modeling of multispecies microbial communities derived directly from metagenomic samples. In short, the pipeline generates metagenome assembled genomes from metagenomic data which are subsequently used to reconstruct and simulate genome scale metabolic models (GEMs) in their communities.

We showed the versatility and usefulness of the metaGEM pipeline by generating metagenome assembled genomes, annotating their taxonomy, calculating relative abundances, reconstructing genome scale models, and simulating metabolic interactions in microbial communities from a range of metagenomic datasets including lab cultures, human gut, plant associated, soil, and ocean metagenomes. Notably, in small metagenomic communities from lab cultures nearly 90% of generated MAGs were of high quality (>90% of genome completeness and <5% contamination), and calculated MAG abundance estimates were highly correlated to marker gene-based estimates (Figure 2a). While metagenomes from more complex communities yielded more MAGs, they pose a more challenging assembly and binning scenario, resulting in a lower percentage of high quality MAGs (Supplementary figure 4). By comparing the generated metabolic models to previously published reference based GEM collections we demonstrated that metaGEMs have a comparable number of reactions and metabolites, despite tending to have fewer genes (Figure 2b), while metaGEM models have significantly less dead end metabolites compared to AGORA, BiGG, and KBase models (Supplementary figure 6). This suggests that some of these reference based reconstructions may contain extraneous metabolic information. Furthermore, by calculating pairwise metabolic distance estimates between models, we show that metaGEMs capture a similar distribution of enzymatic diversity as compared to reference genome based reconstructions (Figure 2c).

We demonstrated that metaGEMs capture a large degree of intraspecies variation by analyzing the core and pangenomes of the top ten most prevalent species from the gut microbiome dataset. Indeed, no two models from the same species were identical in terms of their metabolism, with up to 60% of metabolic diversity was present within species pangenomes demonstrating remarkable degree of intraspecies metabolic variation captured by metaGEM models (Figure 2d). Furthermore, by comparing metaGEMs with reference-based gut species metabolic AGORA models^15^, we showed that reference-based models introduce metabolic reactions that may not necessarily be present in every metagenomic context, while the metaGEM models reconstruct context specific metabolism, entirely based on actual metagenomic data that would have been otherwise missing from the reference-based reconstructions (Figure 2e). Indeed, we find that the most common pathways found exclusively in either metaGEM models and AGORA correspond to the biosynthesis of antibiotics and secondary metabolites, which are known to be context specific and horizontally transferred^61^.

We showed that gut metagenomes corresponding to different type 2 diabetes disease groups (NGT, IGT, T2D)^21^ generate communities with different metabolic architectures. By carrying out species metabolic coupling analysis^17^ for each metagenome-derived personalized community of GEMs, we identified 22 growth-related metabolic exchanges, 27 donor species, and 27 receiver species that were significantly different between at least one disease condition comparison (Supplementary figures 11,13,14). Visualization of SMETANA simulation results revealed notable differences in the metabolic idiosyncrasies of species and communities as a function of their disease state condition, as reflected by scores of their metabolic dependencies (Figure 3). By directly modeling community level sample-specific microbiomes here we identified potential interaction differences between pathogenicity of type 2 diabetes that are based purely on metabolic capacity of the gut communities and are not dependent on species abundance estimates, thus providing additional information that would not be otherwise accessed. Moreover, SMETANA framework is free from arbitrary assumptions of growth optimality, instead, it evaluates all scenarios of interspecies metabolic exchanges that support the growth of member species in a given community providing unbiased community level metabolism analysis^17^. The framework has been validated previously^17^ by reproducing experimentally determined interactions in well-studied microbial communities^62,63^. Overall, our study offers a end-to-end framework to study sample-specific metabolism of complex microbial communities directly from metagenomic data without relying on reference genomes. We therefore envisage that metGEM will become an important tool for deciphering microbial interactions in complex communities.

## Methods

### Tools used by metaGEM

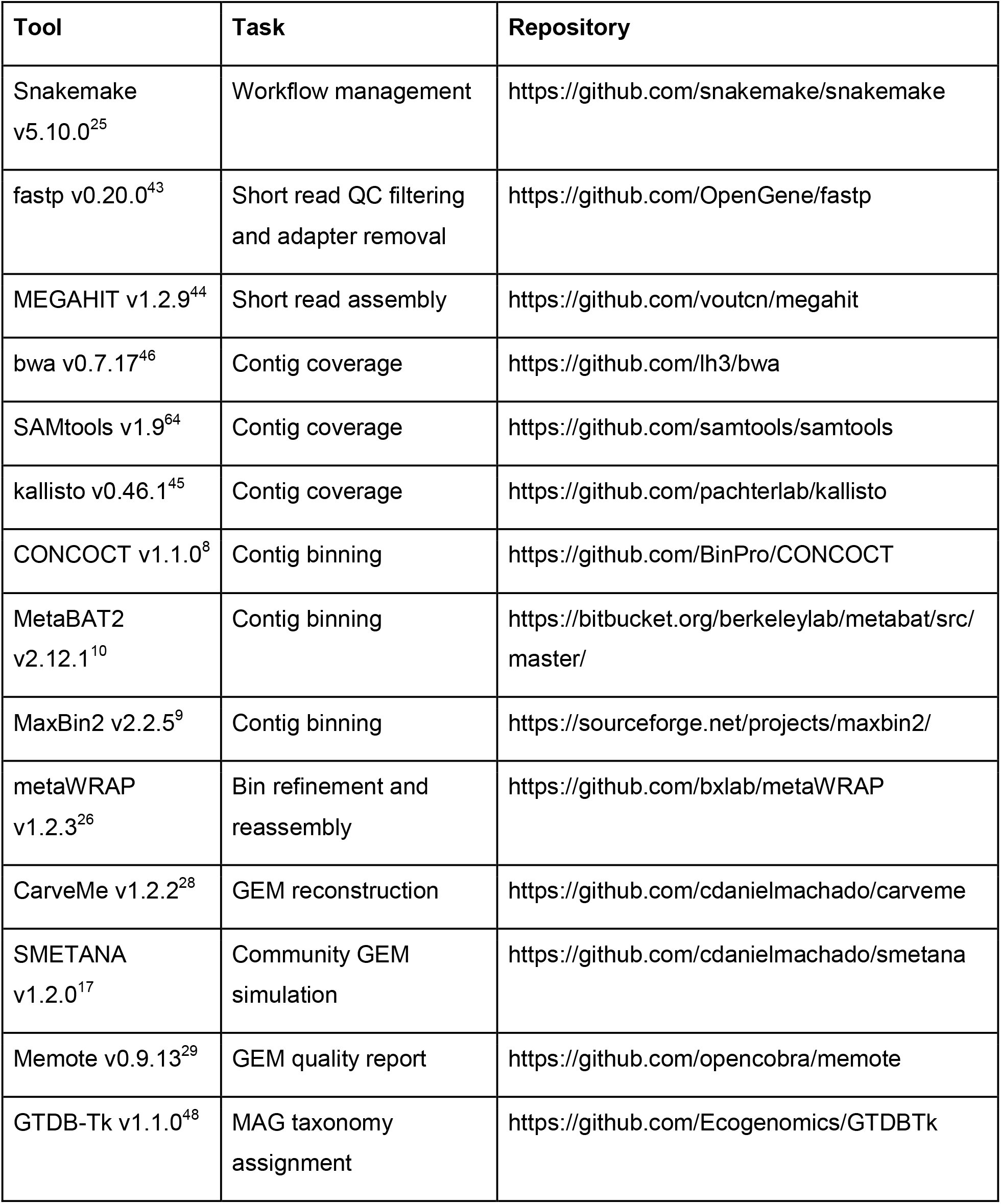
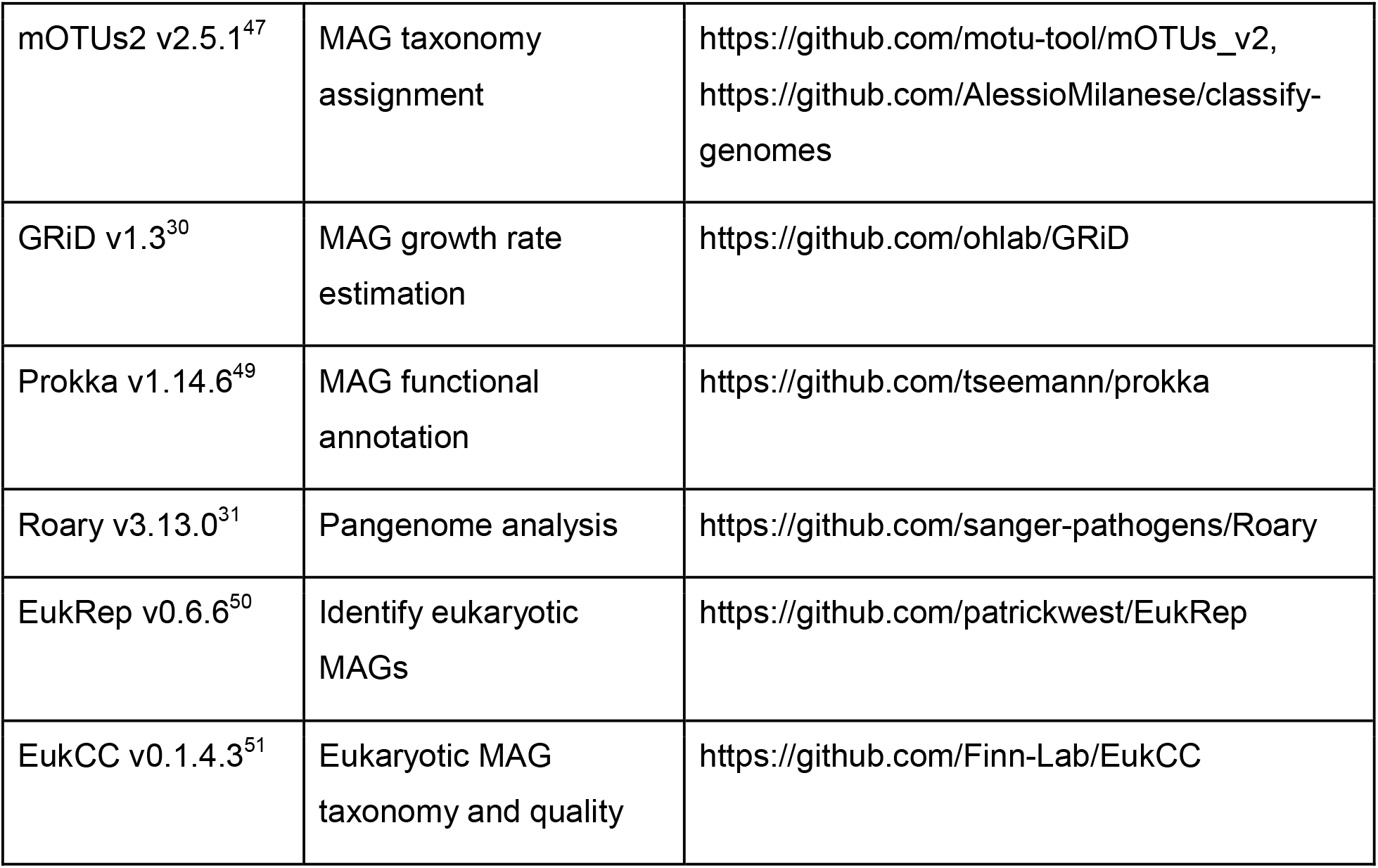

| Tool | Task | Repository |
| --- | --- | --- |
| Snakemake<br>v5.10.0<sup>25</sup> | Workflow management | https://github.com/snakemake/snakemake |
| fastp v0.20.0<sup>43</sup> | Short read QC filtering<br>and adapter removal | https://github.com/OpenGene/fastp |
| MEGAHIT v1.2.9<sup>44</sup> | Short read assembly | https://github.com/voutcn/megahit |
| bwa v0.7.17<sup>46</sup> | Contig coverage | https://github.com/lh3/bwa |
| SAMtools v1.9<sup>64</sup> | Contig coverage | https://github.com/samtools/samtools |
| kallisto v0.46.1<sup>45</sup> | Contig coverage | https://github.com/pachterlab/kallisto |
| CONCOCT v1.1.0<sup>8</sup> | Contig binning | https://github.com/BinPro/CONCOCT |
| MetaBAT2<br>v2.12.1<sup>10</sup> | Contig binning | https://bitbucket.org/berkeleylab/metabat/src/<br>master/ |
| MaxBin2 v2.2.5<sup>9</sup> | Contig binning | https://sourceforge.net/projects/maxbin2/ |
| metaWRAP<br>v1.2.3<sup>26</sup> | Bin refinement and<br>reassembly | https://github.com/bxlab/metaWRAP |
| CarveMe v1.2.2<sup>28</sup> | GEM reconstruction | https://github.com/cdanielmachado/carveme |
| SMETANA<br>v1.2.0<sup>17</sup> | Community GEM<br>simulation | https://github.com/cdanielmachado/smetana |
| Memote v0.9.13<sup>29</sup> | GEM quality report | https://github.com/opencobra/memote |
| GTDB-Tk v1.1.0<sup>48</sup> | MAG taxonomy<br>assignment | https://github.com/Ecogenomics/GTDBTk |
| mOTUs2 v2.5.1<sup>47</sup> | MAG taxonomy assignment | https://github.com/motu-tool/mOTUs_v2,<br>https://github.com/AlessioMilanese/classify-genomes |
| GRiD v1.3<sup>30</sup> | MAG growth rate estimation | https://github.com/ohlab/GRiD |
| Prokka v1.14.6<sup>49</sup> | MAG functional annotation | https://github.com/tseemann/prokka |
| Roary v3.13.0<sup>31</sup> | Pangenome analysis | https://github.com/sanger-pathogens/Roary |
| EukRep v0.6.6<sup>50</sup> | Identify eukaryotic MAGs | https://github.com/patrickwest/EukRep |
| EukCC v0.1.4.3<sup>51</sup> | Eukaryotic MAG taxonomy and quality | https://github.com/Finn-Lab/EukCC |

### Snakemake and HPCCs

metaGEM was implemented in Snakemake v5.10.0, and designed to analyze datasets independently from each other, while samples from the same datasets are processed in parallel. All Snakemake jobs were run on the Chalmers Center for Computational Science and Engineering (C3SE) and the European Molecular Biology Laboratory (EMBL) Heidelberg high performance computer clusters (HPCC).

### Metagenomic samples and short read quality control

A total of 483 whole metagenome shotgun samples from five metagenomic studies^20–24^ were downloaded from the NCBI SRA or EBI ENA to the HPCC. The lab culture dataset was the only single end read set, while the remaining datasets consisted of paired end read sets. The fastp tool v0.20.0 was used for short read quality filtering and adapter removal using default settings.

### Short read assembly

Single sample assemblies were obtained using MEGAHIT v1.2.9. The flag ‘--presets meta-sensitive’ was used on all assemblies, which is equivalent to setting ‘--min-count 1’ ‘--k-list 21,29,39,49,59,69,79,89,99,109,119,129,141’. The parameter ‘--min-contig-len’ was set to 1000 for the TARA Oceans dataset.

### Contig coverage estimation and binning

For the lab culture, gut microbiome, plant associated, and soil datasets, bwa-mem v0.7.17 was used with default settings to cross map each quality controlled set of short reads to each generated assembly within a dataset. An index was created for each assembly prior to mapping using the ‘bwa index’ command with default settings. Each mapping operation resulted in a SAM file, which was converted to BAM format using the ‘samtools view’ command with the flags ‘-Sb’, and then sorted using the ‘samtools sort’ command with default settings. The sorted BAM files were then used to create the contig coverage across samples input files for MetaBAT2 v.2.12.1 and MaxBin2 v2.2.5 with the MetaBAT2 script ‘jgi_summarize_bam_contig_depths’ (default parameters). The CONCOCT script ‘cut_up_fasta.py’ with parameters ‘-c 10000 -o 0 –m -b’ was used to generate a BEDfile for each assembly, with contigs cut into 10 kbp chunks. The generated BEDfile and sorted BAM files were used by the CONCOCT script ‘concoct_coverage_table.py’ (default settings) to generate the contig coverage across samples input files for CONCOCT. CONCOCT v1.1.0 was run for each cut up assembly, using the contig coverage across samples as input and with the ‘-c 800’ parameter. The CONCOCT script ‘extract_fasta_bins.py’ script was then run with default settings to extract bins in terms of the original uncut contigs. MetaBAT2 v.2.12.1 was run using the ‘metabat2’ command for each assembly, using the contig coverage across samples as input and with default settings. MaxBin2 v2.2.5 was run using the ‘run_MaxBin.pl’ for each assembly, using the contig coverage across samples as input and with default settings.

For the TARA oceans dataset, the kallisto v0.46.1 tool was used for mapping quality controlled paired end reads to assemblies. First, each assembly was cut up into 10 kbp chunks using the CONCOCT ‘cut_up_fasta.py ‘ script with parameters ‘-c 10000 -o 0 -m’. The cut up assembly was then used to generate a kallisto index using the ‘kallisto index’ command with default settings. Next, the ‘kallisto quant’ command was used with the ‘--plaintext’ setting to cross map each sample set of quality controlled paired end reads to each cut up assembly. Finally, the ‘kallisto2concoct.py’ script was used to summarize the mapping results across samples for each set of assembled contigs. For the TARA oceans dataset the binners were used as described above, but only CONCOCT used contig coverage across samples as input, while MetaBAT2 and MaxBin2 only used contig coverage from the assembled sample as input.

### Bin refinement and reassembly

To reconcile and dereplicate the three generated binner outputs, the metaWRAP metaWRAP v1.2.3 ‘bin_refinement’ command was used with parameters ‘-x 10 -c 50’, corresponding to a maximum bin contamination threshold of 10% and a minimum bin completeness threshold of 50% (i.e. medium quality bin criteria) based on CheckM estimates. The metaWRAP ‘reassemble_bins’ command was used with parameters ‘-x 10 -c 50’ to improve bin quality whenever possible by independently reassembling reads that map from the parent sample’s quality controlled short reads to the refined bin. Refined and reassembled bins are hereafter referred to as metagenome assembled genomes (MAGs), although the terms may be used interchangeably.

### MAG abundance quantification and taxonomic assignment

Absolute and relative abundances of MAGs were calculated using bwa v0.7.17 and SAMtools v1.9. First, all MAGs generated from the same sample were concatenated into a single fasta file, based on which an index was created using ‘bwa index’ (default parameters). Next, ‘bwa-mem’ (default parameters) was used to map the quality controlled short reads from the parent sample to the concatenation of generated MAGs. The resulting SAM file was converted to BAM format and sorted using ‘samtools view’ and ‘samtools sort’, respectively, using the same parameters as described above. Next, ‘samtools flagstat’ was used (default parameters) to extract the number of reads that mapped from the quality controlled paired end reads to the concatenation of all MAGs generated in that sample. Next, for each MAG, an index was created using ‘bwa index’ (default parameters), the quality controlled short reads were mapped to the MAG using ‘bwa-mem’ (default parameters), the resulting SAM file was converted to BAM format and sorted as described above, and the number of reads mapping to the MAG from the short reads was extracted using ‘samtools flagstat’. The abundance of a given MAG was estimated by dividing the number of quality controlled reads that map to the MAG by the number of quality controlled reads that map to the concatenation of all MAGs from that sample, divided by the megabase pair length of the MAG. For each sample, these non-normalized abundances were summed to obtain a sample specific normalizing factor. To obtain normalized relative abundances, each non-normalized abundance was divided by the sample specific normalization factor. The mOTUs2 v2.5.1 tool was also used to calculate abundances, for comparison with the above mapping based method, only in the lab culture dataset. The ‘motus profile’ command was used with default settings. GTDB-tk v1.1.0 was used to assign taxonomic labels to the generated MAGs using the ‘gtdbtk classify_wf’ with default settings.

### Genome scale metabolic model reconstruction and quality reports

CarveMe v1.2.2 was used to automatically reconstruct genome scale metabolic models from ORF annotated protein fasta files derived from MAGs using the default CPLEX solver. The ‘carve’ command was run using the ‘--fbc2 -g’ flags to gapfill models on complete media and generate FBC2 format models. The memote v0.9.13 tool was then used to generate quality reports for each genome scale metabolic model. The ‘memote run’ command was used with the flags ‘--skip test_find_metabolites_produced_with_closed_bounds --skip test_find_metabolites_consumed_with_closed_bounds --skip test_find_metabolites_not_produced_with_open_bounds --skip test_find_metabolites_not_consumed_with_open_bounds --skip test_find_incorrect_thermodynamic_reversibility’ to avoid running time consuming tests.

### Simulations

The SMETANA v1.2.0 tool was used for simulating gut microbiome communities of reconstructed genome scale metabolic models. The ‘smetana’ command was used with the flags ‘--flavor fbc2 --detailed --mediadb media_db.tsv -m M11’ and using the default CPLEX solver. The simulation media was the same as was used for gapfilling (full media, M3) minus aromatic amino acids (M11). The media file was obtained from the authors of previous publication^16^, and can be accessed from the metaGEM GitHub repository (https://github.com/franciscozorrilla/metaGEM).

### Additional features in metaGEM pipeline

Although not discussed in detail, there are several additional features that were incorporated into metaGEM that may be useful to users. Growth rates for medium and high coverage MAGs can be estimated using the GRiD v1.3 tool. Prokka v1.14.6 can be used to functionally annotate MAGs and the output can be provided to Roary v3.13.0 in order to visualize the core and pangenome of a set of MAGs. Communities with suspected eukaryotic MAGs can be further probed by scanning for eukaryotic contigs in the CONCOCT bin set using EukRep v0.6.6. Identified eukaryotic bins can then be analyzed by EukCC v0.1.4.3, to obtain completeness and contamination estimates as well taxonomic lineage estimates.

## Data availability

All simulations-ready metagenome-based metabolic reconstructions were deposited to the Zenodo repository and are available at http://doi.org/10.5281/zenodo.4407746.

## Code availability

The code of metaGEM can be accessed at https://github.com/franciscozorrilla/metaGEM.

## Author contributions

FZ, AZ, KRP conceptualized the project; FZ and AZ designed the computational analysis; FZ performed the computational analysis; FZ and AZ interpreted the results; FZ, AZ wrote the initial draft manuscript; FZ, KRP, and AZ revised the draft and wrote the final manuscript.

## Acknowledgements

This study was funded by the DD-DeCaF consortium European Union’s Horizon 2020 research and innovation programme [686070]. AZ was funded by the SciLifeLab fellows program. The computations were enabled with resources provided by EMBL and by the Swedish National Infrastructure for Computing (SNIC) at C3SE partially funded by the Swedish Research Council through grant agreement no. 2018-05973. Mikael Öhman and Thomas Svedberg at C3SE are acknowledged for technical assistance in making the code run on Vera C3SE resources.

## References

1. Quince, C. et al. DESMAN: a new tool for de novo extraction of strains from metagenomes. Genome Biol. 18, 181 (2017).

2. Durack, J. & Lynch, S. V. The gut microbiome: Relationships with disease and opportunities for therapy. J. Exp. Med. 216, 20–40 (2019).

3. Sanna, S. et al. Causal relationships among the gut microbiome, short-chain fatty acids and metabolic diseases. Nat. Genet. 51, 600–605 (2019).

4. Li, Q. et al. Implication of the gut microbiome composition of type 2 diabetic patients from northern China. Sci. Rep. 10, 5450 (2020).

5. Gopalakrishnan, V., Helmink, B. A., Spencer, C. N., Reuben, A. & Wargo, J. A. The Influence of the Gut Microbiome on Cancer, Immunity, and Cancer Immunotherapy. Cancer Cell 33, 570–580 (2018).

6. Vuong, H. E. & Hsiao, E. Y. Emerging Roles for the Gut Microbiome in Autism Spectrum Disorder. Biol. Psychiatry 81, 411–423 (2017).

7. Cryan, J. F., O’Riordan, K. J., Sandhu, K., Peterson, V. & Dinan, T. G. The gut microbiome in neurological disorders. Lancet Neurol. 19, 179–194 (2020).

8. Alneberg, J. et al. Binning metagenomic contigs by coverage and composition. Nat. Methods 11, 1144–1146 (2014).

9. Wu, Y.-W., Simmons, B. A. & Singer, S. W. MaxBin 2.0: an automated binning algorithm to recover genomes from multiple metagenomic datasets. Bioinformatics 32, 605–607 (2015).

10. Kang, D. D. et al. MetaBAT 2: an adaptive binning algorithm for robust and efficient genome reconstruction from metagenome assemblies. PeerJ 7, e7359 (2019).

11. Almeida, A. et al. A new genomic blueprint of the human gut microbiota. Nature 568, 499–504 (2019).

12. Nayfach, S., Shi, Z. J., Seshadri, R., Pollard, K. S. & Kyrpides, N. C. New insights from uncultivated genomes of the global human gut microbiome. Nature 568, 505–510 (2019).

13. Pasolli, E. et al. Extensive Unexplored Human Microbiome Diversity Revealed by Over 150,000 Genomes from Metagenomes Spanning Age, Geography, and Lifestyle. Cell 176, 649–662.e20 (2019).

14. Almeida, A. et al. A unified catalog of 204,938 reference genomes from the human gut microbiome. Nat. Biotechnol. (2020) doi:10.1038/s41587-020-0603-3.

15. Magnúsdóttir, S. et al. Generation of genome-scale metabolic reconstructions for 773 members of the human gut microbiota. Nat. Biotechnol. 35, 81–89 (2017).

16. Tramontano, M. et al. Nutritional preferences of human gut bacteria reveal their metabolic idiosyncrasies. Nat Microbiol 3, 514–522 (2018).

17. Zelezniak, A. et al. Metabolic dependencies drive species co-occurrence in diverse microbial communities. Proceedings of the National Academy of Sciences 112, 6449 LP–6454 (2015).

18. Basile, A. et al. Revealing metabolic mechanisms of interaction in the anaerobic digestion microbiome by flux balance analysis. Metab. Eng. 62, 138–149 (2020).

19. Freilich, S. et al. Competitive and cooperative metabolic interactions in bacterial communities. Nat. Commun. 2, 589 (2011).

20. Korem, T. et al. Growth dynamics of gut microbiota in health and disease inferred from single metagenomic samples. Science 349, 1101 LP–1106 (2015).

21. Karlsson, F. H. et al. Gut metagenome in European women with normal, impaired and diabetic glucose control. Nature 498, 99–103 (2013).

22. Li, X. et al. Legacy of land use history determines reprogramming of plant physiology by soil microbiome. ISME J. 13, 738–751 (2019).

23. Sunagawa, S. et al. Structure and function of the global ocean microbiome. Science 348, 1261359 (2015).

24. Bissett, A. et al. Introducing BASE: the Biomes of Australian Soil Environments soil microbial diversity database. Gigascience 5, 21 (2016).

25. Köster, J. & Rahmann, S. Snakemake—a scalable bioinformatics workflow engine. Bioinformatics 28, 2520–2522 (2012).

26. Uritskiy, G. V., DiRuggiero, J. & Taylor, J. MetaWRAP—a flexible pipeline for genome-resolved metagenomic data analysis. Microbiome 6, 158 (2018).

27. Parks, D. H., Imelfort, M., Skennerton, C. T., Hugenholtz, P. & Tyson, G. W. CheckM: assessing the quality of microbial genomes recovered from isolates, single cells, and metagenomes. Genome Res. 25, 1043–1055 (2015).

28. Machado, D., Andrejev, S., Tramontano, M. & Patil, K. R. Fast automated reconstruction of genome-scale metabolic models for microbial species and communities. Nucleic Acids Res. 46, 7542–7553 (2018).

29. Lieven, C. et al. MEMOTE for standardized genome-scale metabolic model testing. Nat. Biotechnol. 38, 272–276 (2020).

30. Emiola, A. & Oh, J. High throughput in situ metagenomic measurement of bacterial replication at ultra-low sequencing coverage. Nat. Commun. 9, 4956 (2018).

31. Page, A. J. et al. Roary: rapid large-scale prokaryote pan genome analysis. Bioinformatics 31, 3691–3693 (2015).

32. Murat Eren, A. et al. Anvi’o: an advanced analysis and visualization platform for ‘omics data. PeerJ 3, e1319 (2015).

33. Goecks, J., Nekrutenko, A., Taylor, J. & The Galaxy Team. Galaxy: a comprehensive approach for supporting accessible, reproducible, and transparent computational research in the life sciences. Genome Biol. 11, R86 (2010).

34. Tamames, J. & Puente-Sánchez, F. SqueezeMeta, A Highly Portable, Fully Automatic Metagenomic Analysis Pipeline. Frontiers in Microbiology vol. 9 3349 (2019).

35. Clarke, E. L. et al. Sunbeam: an extensible pipeline for analyzing metagenomic sequencing experiments. Microbiome 7, 46 (2019).

36. Murovec, B., Deutsch, L. & Stres, B. Computational Framework for High-Quality Production and Large-Scale Evolutionary Analysis of Metagenome Assembled Genomes. Mol. Biol. Evol. 37, 593–598 (2019).

37. Stewart, R. D., Auffret, M. D., Snelling, T. J., Roehe, R. & Watson, M. MAGpy: a reproducible pipeline for the downstream analysis of metagenome-assembled genomes (MAGs). Bioinformatics 35, 2150–2152 (2018).

38. Arkin, A. P. et al. KBase: The United States Department of Energy Systems Biology Knowledgebase. Nat. Biotechnol. 36, 566–569 (2018).

39. Tully, B. J., Sachdeva, R., Graham, E. D. & Heidelberg, J. F. 290 metagenome-assembled genomes from the Mediterranean Sea: a resource for marine microbiology. PeerJ 5, e3558–e3558 (2017).

40. Delmont, T. O. et al. Nitrogen-fixing populations of Planctomycetes and Proteobacteria are abundant in surface ocean metagenomes. Nature Microbiology 3, 804–813 (2018).

41. Tully, B. J., Graham, E. D. & Heidelberg, J. F. The reconstruction of 2,631 draft metagenome-assembled genomes from the global oceans. Scientific Data 5, 170203 (2018).

42. Zrimec, J., Kokina, M., Jonasson, S., Zorrilla, F. & Zelezniak, A. Plastic-degrading potential across the global microbiome correlates with recent pollution trends. doi:10.1101/2020.12.13.422558.

43. Chen, S., Zhou, Y., Chen, Y. & Gu, J. fastp: an ultra-fast all-in-one FASTQ preprocessor. Bioinformatics 34, i884–i890 (2018).

44. Li, D., Liu, C.-M., Luo, R., Sadakane, K. & Lam, T.-W. MEGAHIT: an ultra-fast single-node solution for large and complex metagenomics assembly via succinct de Bruijn graph. Bioinformatics 31, 1674–1676 (2015).

45. Bray, N. L., Pimentel, H., Melsted, P. & Pachter, L. Near-optimal probabilistic RNA-seq quantification. Nat. Biotechnol. 34, 525–527 (2016).

46. Li, H. Aligning sequence reads, clone sequences and assembly contigs with BWA-MEM. arXiv [q-bio.GN] (2013).

47. Milanese, A. et al. Microbial abundance, activity and population genomic profiling with mOTUs2. Nat. Commun. 10, 1014 (2019).

48. Chaumeil, P.-A., Mussig, A. J., Hugenholtz, P. & Parks, D. H. GTDB-Tk: a toolkit to classify genomes with the Genome Taxonomy Database. Bioinformatics 36, 1925–1927 (2019).

49. Seemann, T. Prokka: rapid prokaryotic genome annotation. Bioinformatics 30, 2068–2069 (2014).

50. West, P. T., Probst, A. J., Grigoriev, I. V., Thomas, B. C. & Banfield, J. F. Genome-reconstruction for eukaryotes from complex natural microbial communities. Genome Res. 28, 569–580 (2018).

51. Saary, P., Mitchell, A. L. & Finn, R. D. Estimating the quality of eukaryotic genomes recovered from metagenomic analysis. bioRxiv 2019.12.19.882753 (2020).

52. King, Z. A. et al. BiGG Models: A platform for integrating, standardizing and sharing genome-scale models. Nucleic Acids Res. 44, D515–D522 (2015).

53. Zhu, A., Sunagawa, S., Mende, D. R. & Bork, P. Inter-individual differences in the gene content of human gut bacterial species. Genome Biol. 16, 82 (2015).

54. Livingstone, P. G., Morphew, R. M. & Whitworth, D. E. Genome Sequencing and Pan-Genome Analysis of 23 Corallococcus spp. Strains Reveal Unexpected Diversity, With Particular Plasticity of Predatory Gene Sets. Front. Microbiol. 9, 3187 (2018).

55. Gurung, M. et al. Role of gut microbiota in type 2 diabetes pathophysiology. EBioMedicine 51, (2020).

56. Garud, N. R. & Pollard, K. S. Population Genetics in the Human Microbiome. Trends Genet. 36, 53–67 (2020).

57. Peña-Gonzalez, A. et al. Metagenomic Signatures of Gut Infections Caused by Different Escherichia coli Pathotypes. Appl. Environ. Microbiol. 85, (2019).

58. Frioux, C., Singh, D., Korcsmaros, T. & Hildebrand, F. From bag-of-genes to bag-of-genomes: metabolic modelling of communities in the era of metagenome-assembled genomes. Comput. Struct. Biotechnol. J. 18, 1722–1734 (2020).

59. Zou, Y. et al. 1,520 reference genomes from cultivated human gut bacteria enable functional microbiome analyses. Nat. Biotechnol. 37, 179–185 (2019).

60. Kim, C. Y. et al. Human reference gut microbiome comprising 5,414 prokaryotic species, including newly assembled genomes from under-represented Asian metagenomes. doi:10.1101/2020.11.09.375873.

61. Versluis, D. et al. Mining microbial metatranscriptomes for expression of antibiotic 1 resistance genes under natural conditions. Sci. Rep. 5, 11981 (2015).

62. Hom, E. F. Y. & Murray, A. W. Niche engineering demonstrates a latent capacity for fungal-algal mutualism. Science 345, 94–98 (2014).

63. Miller, L. D. et al. Establishment and metabolic analysis of a model microbial community for understanding trophic and electron accepting interactions of subsurface anaerobic environments. BMC Microbiol. 10, 149 (2010).

64. Li, H. et al. The Sequence Alignment/Map format and SAMtools. Bioinformatics 25, 2078–2079 (2009).

