## Supplementary Information for "metaGEM: reconstruction of genome scale metabolic models directly from metagenomes"

### Supplementary Figures

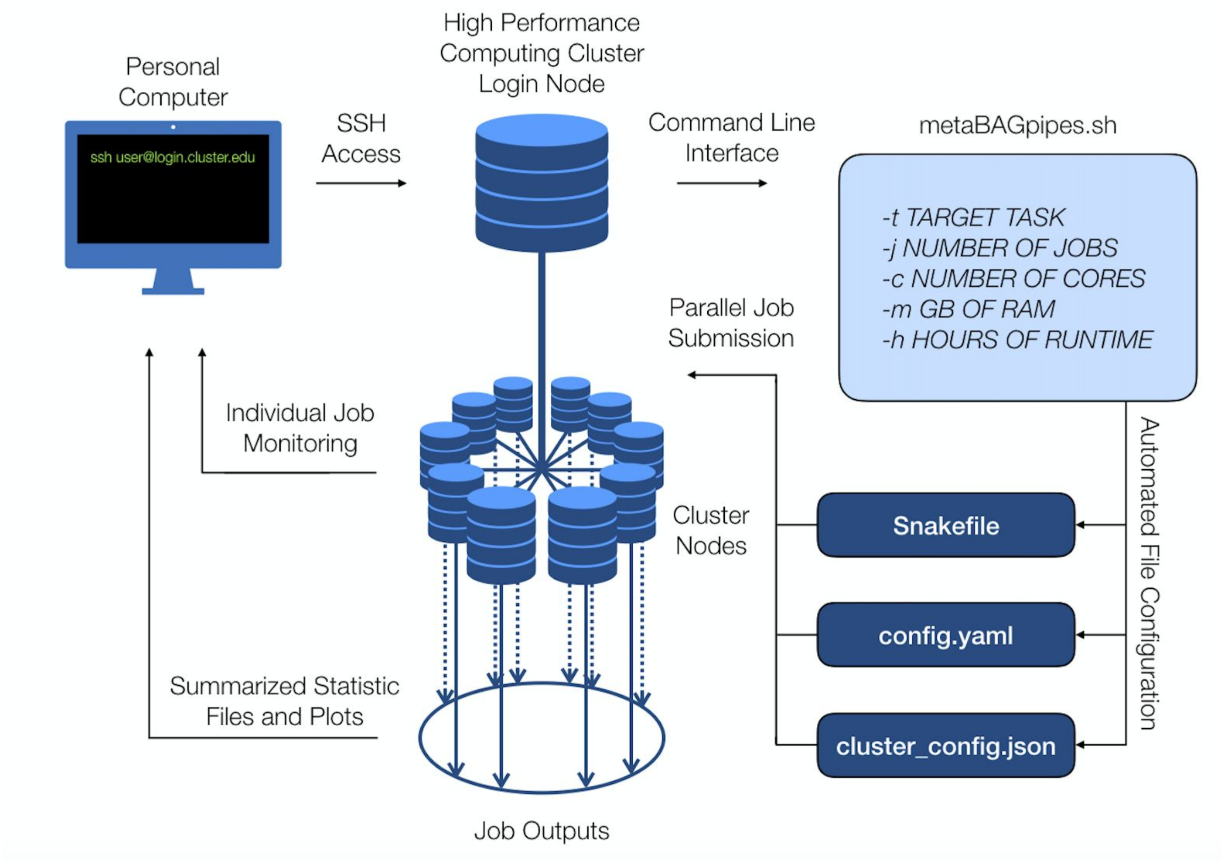


**Supplementary figure 1.** Schematic overview of user interface with high performance computing cluster environment and metaGEM.

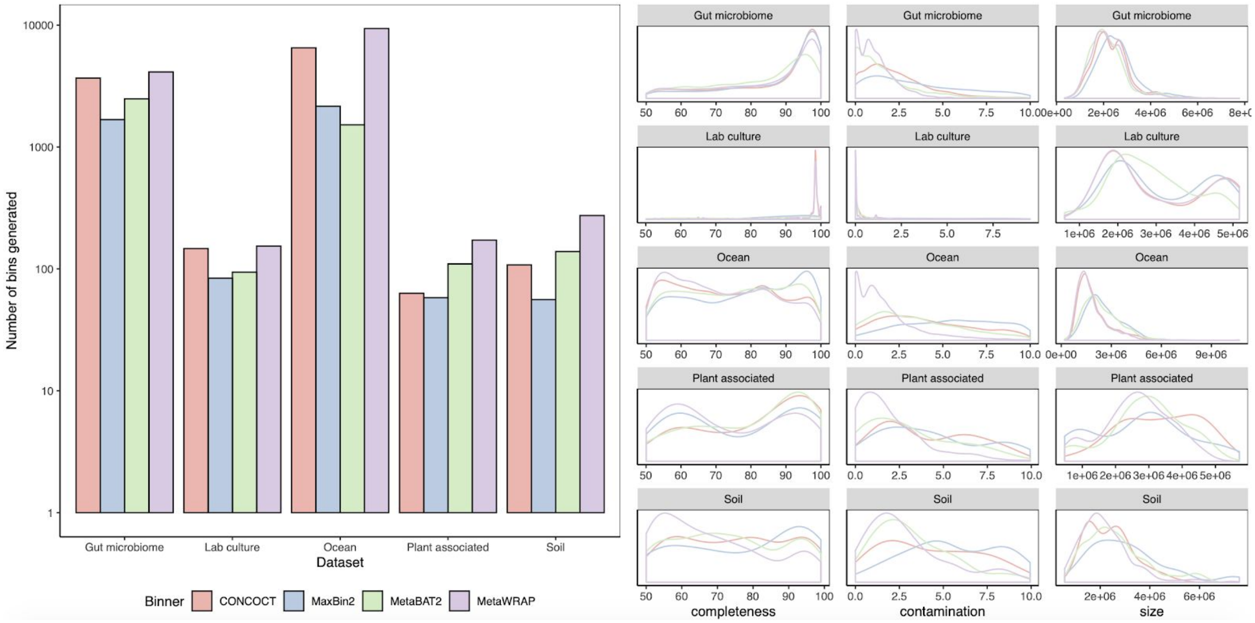


**Supplementary figure 2.** Summary statistics of intermediate binning results, showing that metaWRAP generates the most low contamination bins compared to individual binners. Left panel shows bar plots representing the number of bins generated by CONCOCT, MaxBin2, MetaBAT2, and metaWRAP across datasets. Right panel shows distributions of completeness, contamination, and bin size across datasets for each binner.

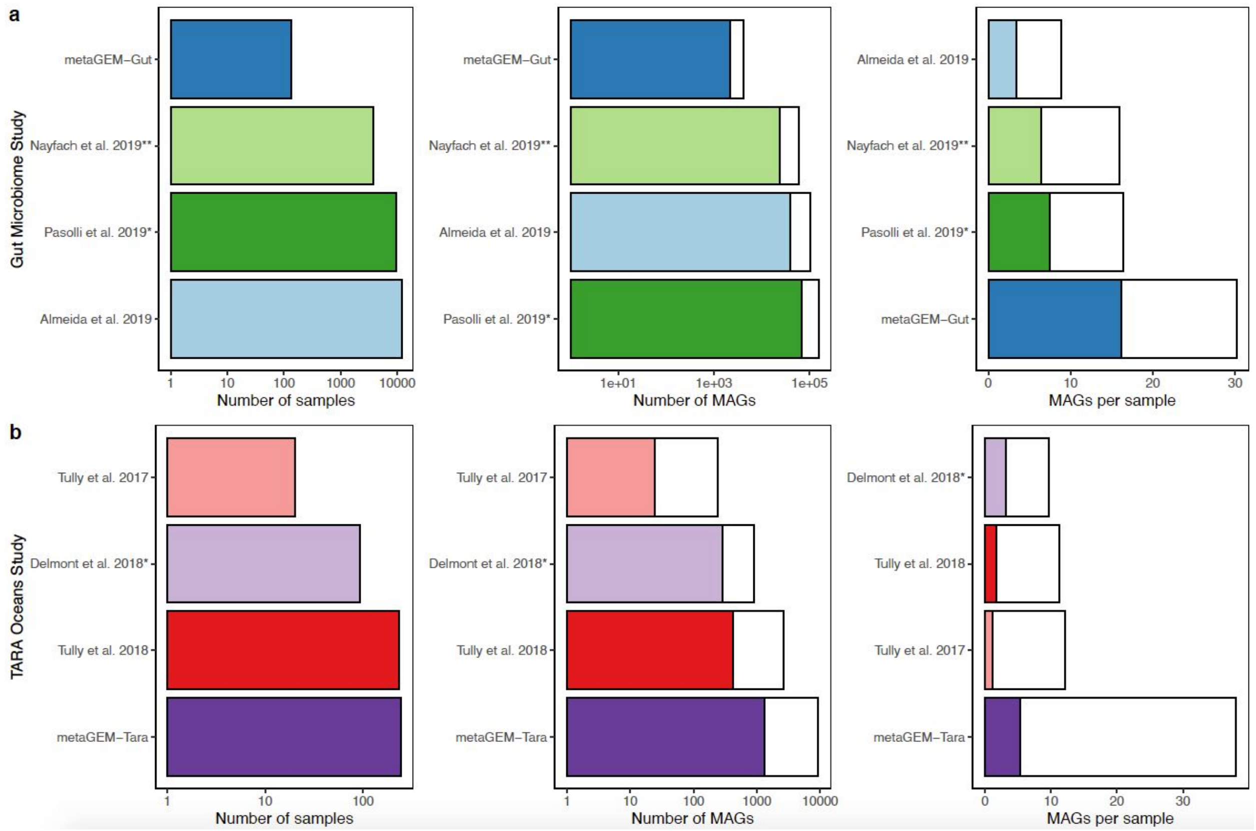


**Supplementary figure 3.** Comparison to other studies showing number of samples used, number of MAGs reconstructed, and number of MAGs reconstructed per sample. (a) Comparisons to large scale gut microbiome MAG reconstruction papers. (b) Comparison to Tara Oceans MAG reconstruction papers. Panel A shows that on average metaGEM reconstructed 30.2 MQ MAGs per sample, 16.1 of which are also HQ. metaGEM generates approximately twice the number of MQ MAGs per sample compared to the methods used in three large scale gut microbiome MAG reconstruction papers. This improved number likely has to do with the cross mapping implementation which is not used by other studies. Panel B shows that the analysis performed by the metaGEM implementation on the Tara Oceans dataset outperforms other approaches. The Tully et al. 2017 study manually assigned taxonomy to each of the 290 reconstructed MAGs, while metaGEM can assign taxonomy automatically. Additionally, only 243 out the 290 reconstructed genomes met the medium quality criteria of >50% completion and <10% contamination. The Delmont et al. 2018 study used an extensive manual curation approach for each reconstructed MAG using the anvi'o workflow, whereas metaGEM requires no manual curation. However, only 899 out of the 1077 redundant MAGs met the medium quality criteria of >50% completion and <10% contamination. Additionally, their approach included a co-assembly approach based on the location of samples. While this may improve MAG assembly quality, it does so at the risk of generating composite or chimeric genomes that do not reflect true biological organisms. While the Tully et al. 2018 paper claimed 603 near complete high quality genomes, only 403 of these passed the >90% completeness and <10% contamination criteria for high quality. metaGEM generates approximately 4 times the number of

MQ MAGs per sample and almost 2 the number of HQ MAGs per sample compared to other methods used to analyse samples from the Tara Oceans bacterial fraction dataset.

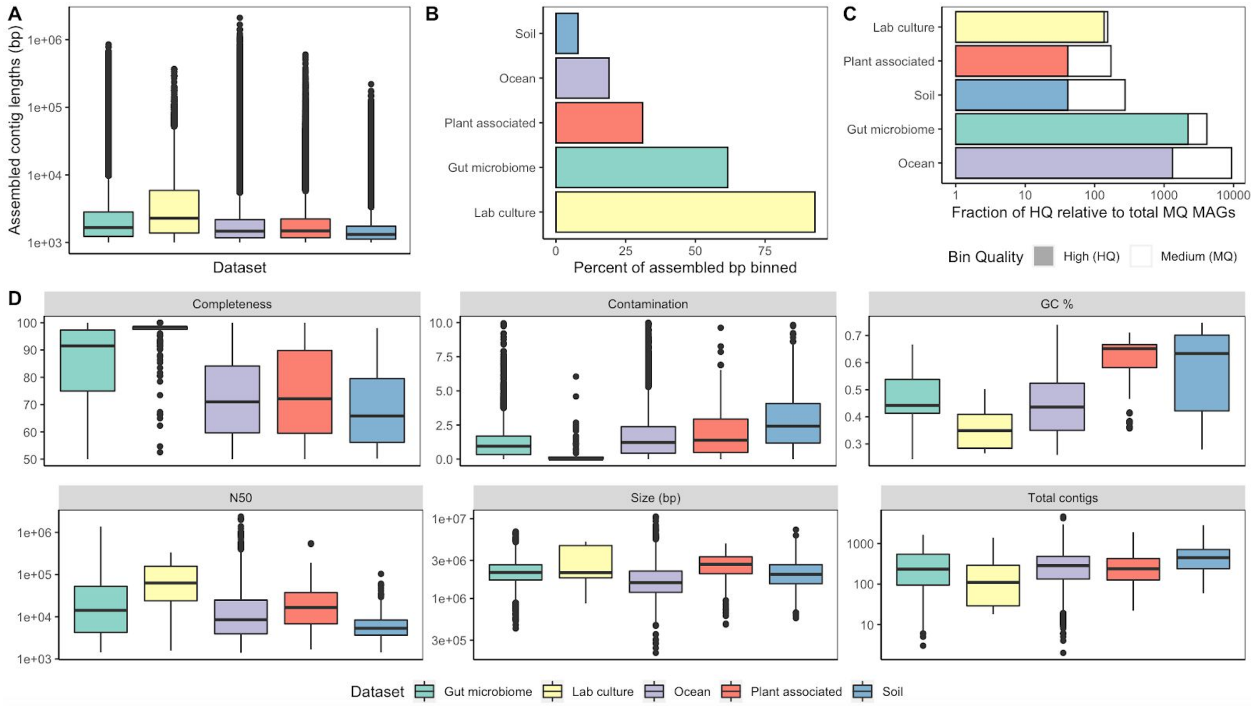


**Supplementary figure 4.** Summarized statistics of assembly and binning process. (A) Boxplots of contig length for all assembled contigs across datasets. (B) Bar plots showing percent of all assembled contigs that end in up at least medium quality bins. (C) Bar plots showing total reconstructed bins across datasets, highlighting the fraction of bins that are also high quality. (D) Distributions of completeness, contamination, GC %, N50, size (bp), and number of contigs across all 14099 reconstructed MQ prokaryotic MAGs.

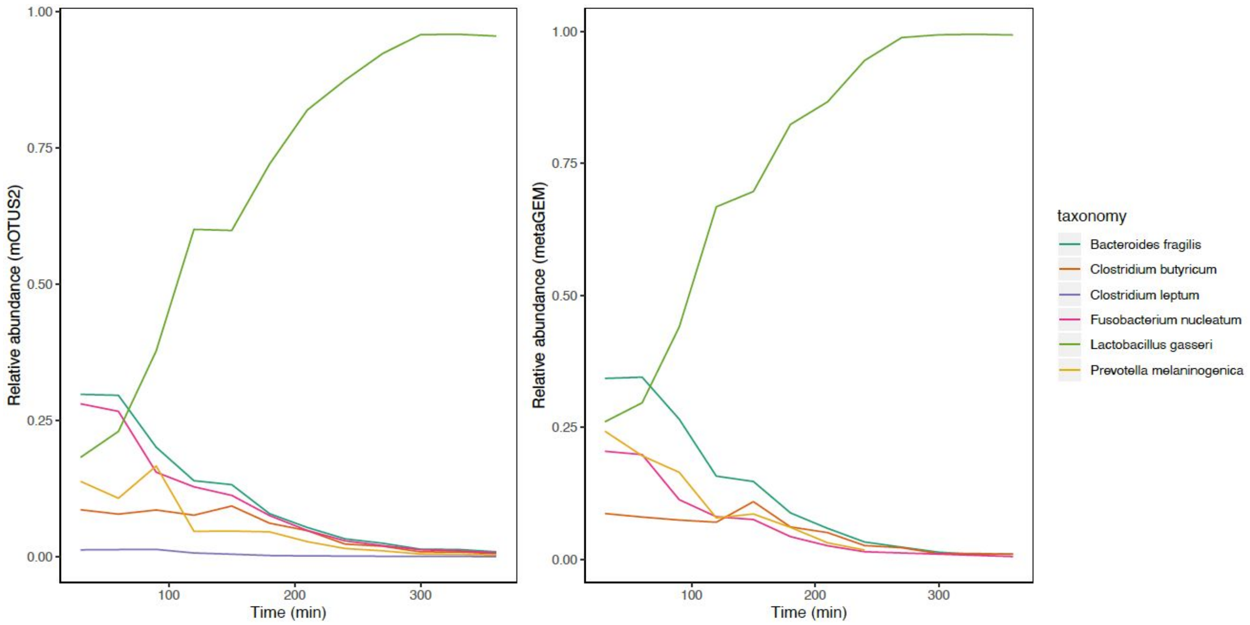


**Supplementary figure 5.** Reconstructed time-course abundance curves from lab culture dataset. Left hand panel shows relative abundances calculated using marker gene based approach (mOTUs2). Right hand panel shows relative abundances calculated using a mapping based approach (metaGEM).

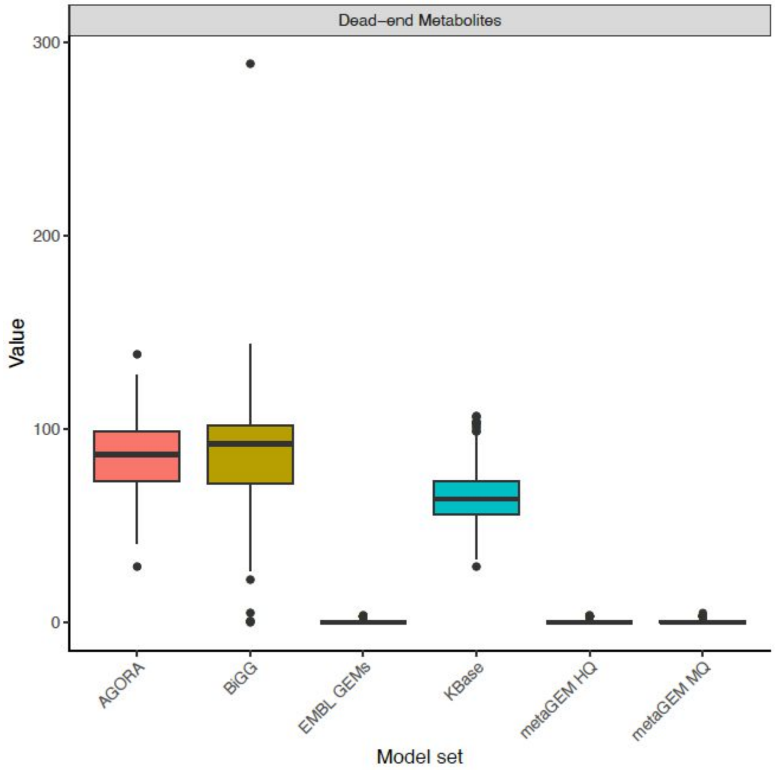


**Supplementary figure 6.** Distributions of number of dead-end metabolites across model sets based on memote reports, with metaGEM models split into medium quality and high quality categories.

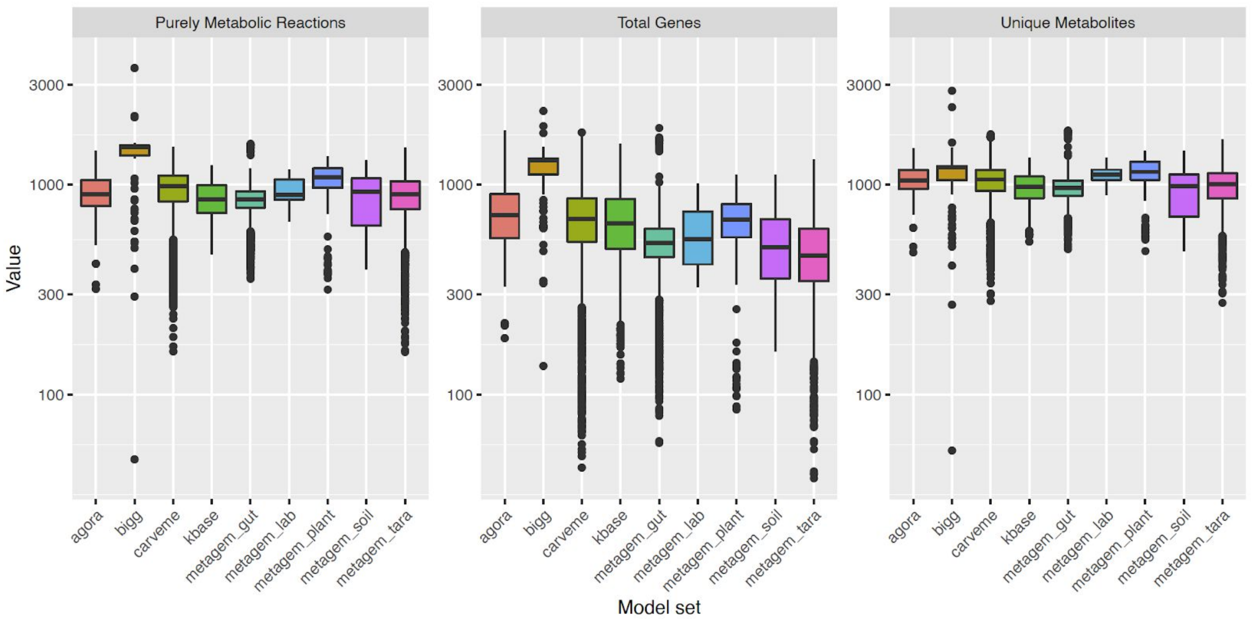


**Supplementary figure 7.** Comparison of key metrics between previously published GEM collections and metaGEM generated models. Left-most panel shows distribution of metabolic reactions in models. Middle panel distribution shows distribution of genes in models. Right-most panel shows distribution of unique metabolites in models.

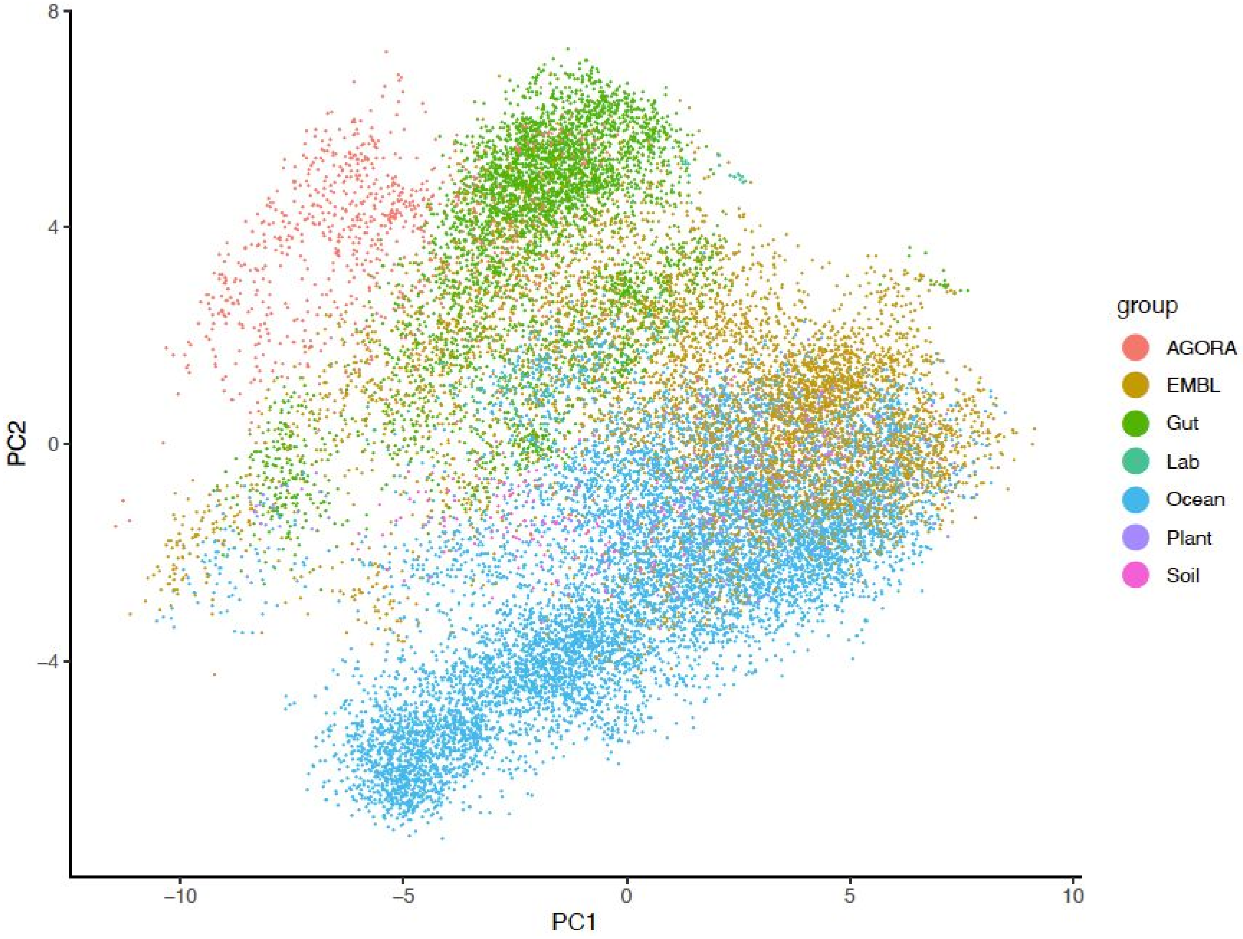


**Supplementary figure 8.** Principal component analysis of reconstructed genome scale metabolic models based on the presence/absence of EC numbers.

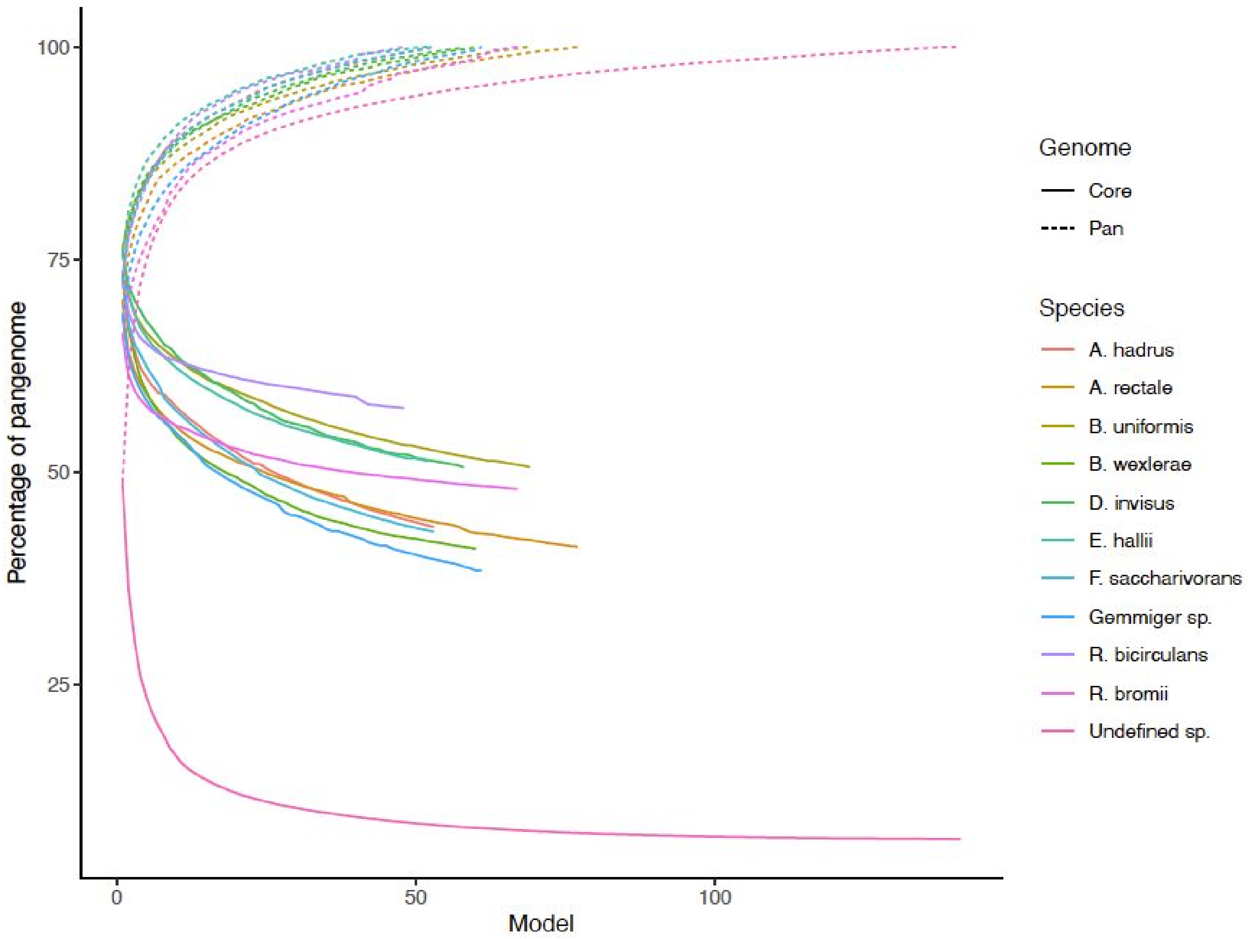


**Supplementary figure 9.** Cumulative core and pan genome curves for the top eleven most commonly reconstructed gut microbiome species, including undefined species, based on EC numbers present in the reconstructed genome scale metabolic models.

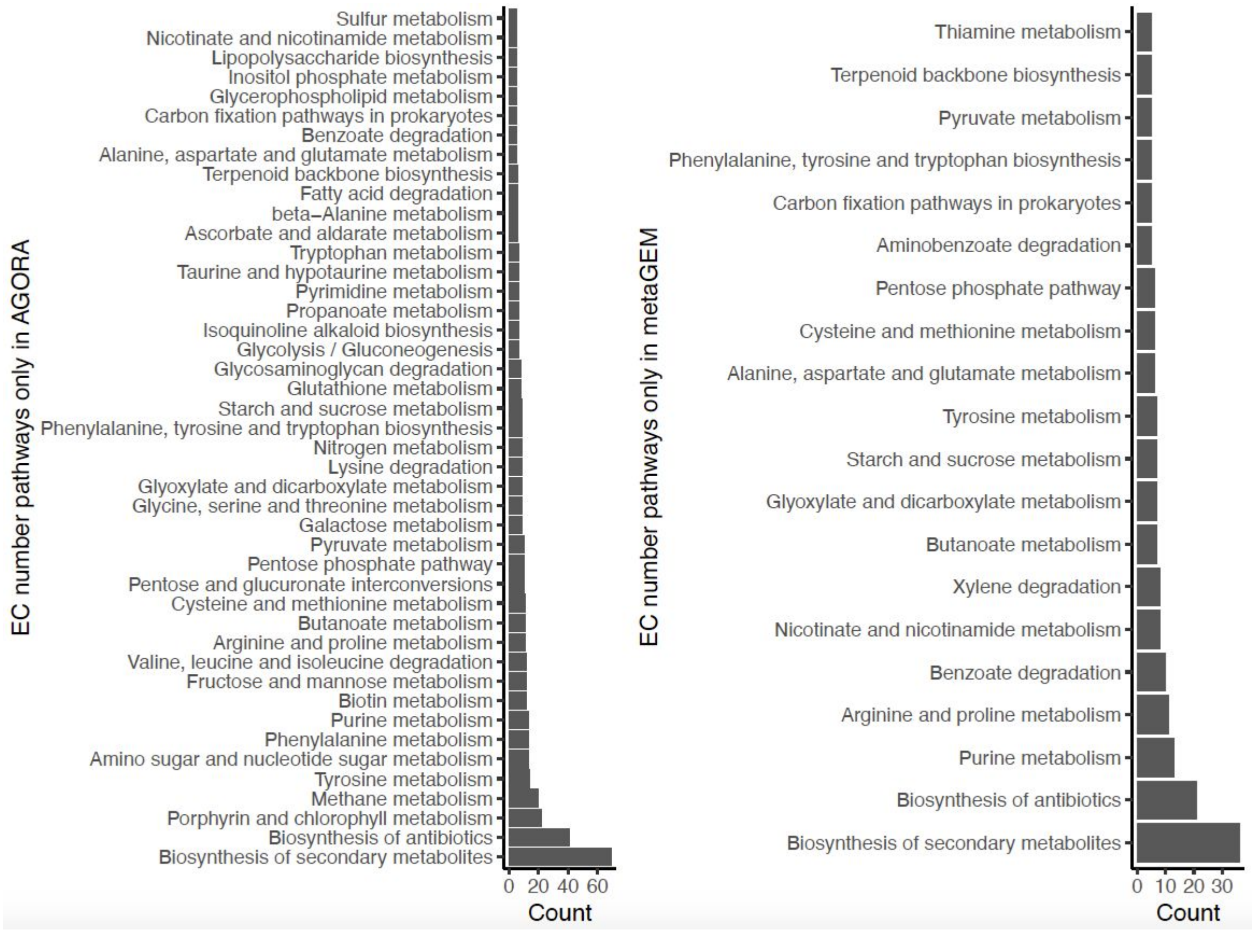


**Supplementary figure 10.** Pathways present only in AGORA model and pathways present only in metaGEM model based on EC number comparison of 165 matched species.

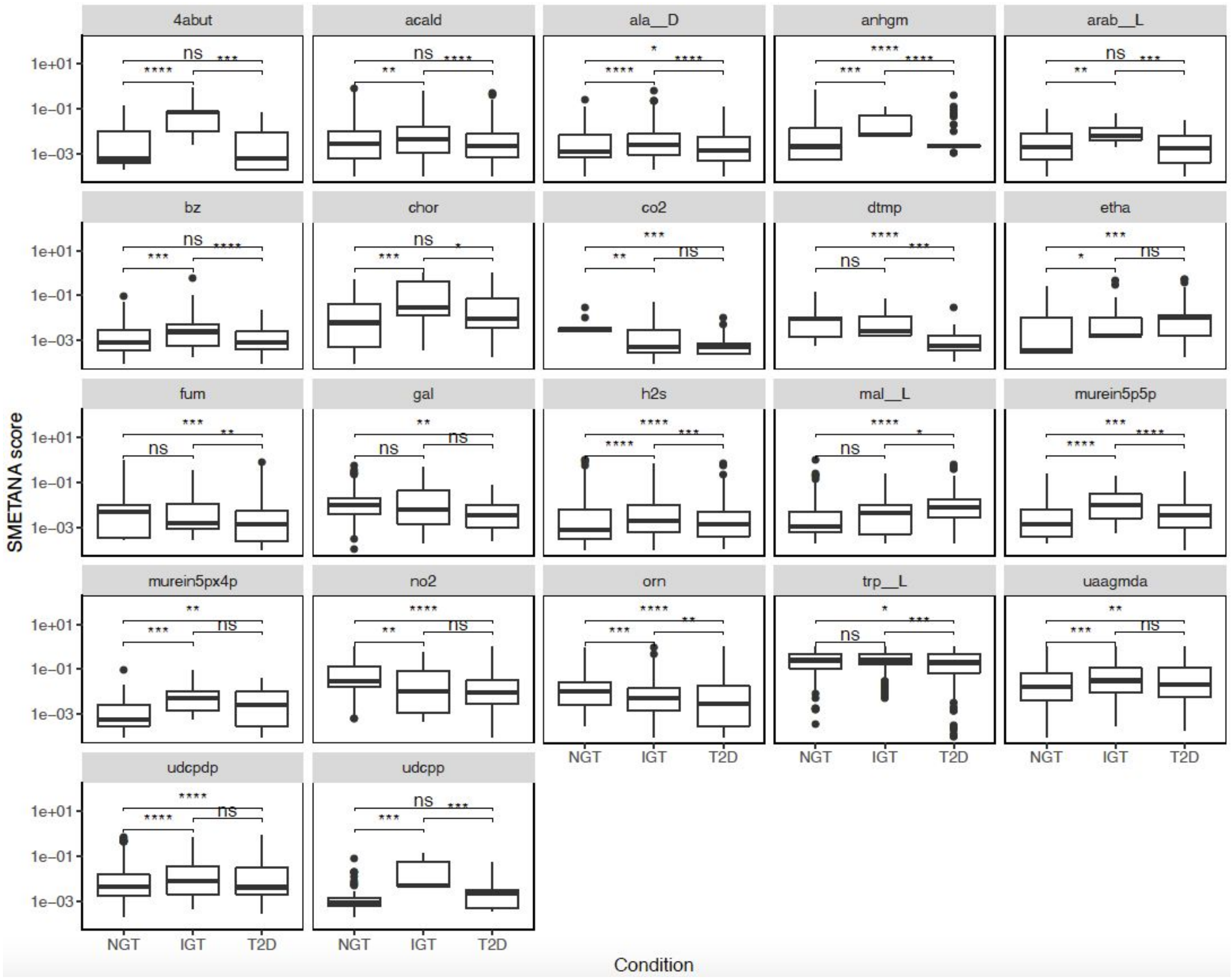


**Supplementary figure 11.** Distribution of SMETNA scores for 22 metabolites with statistically significant differences across conditions.

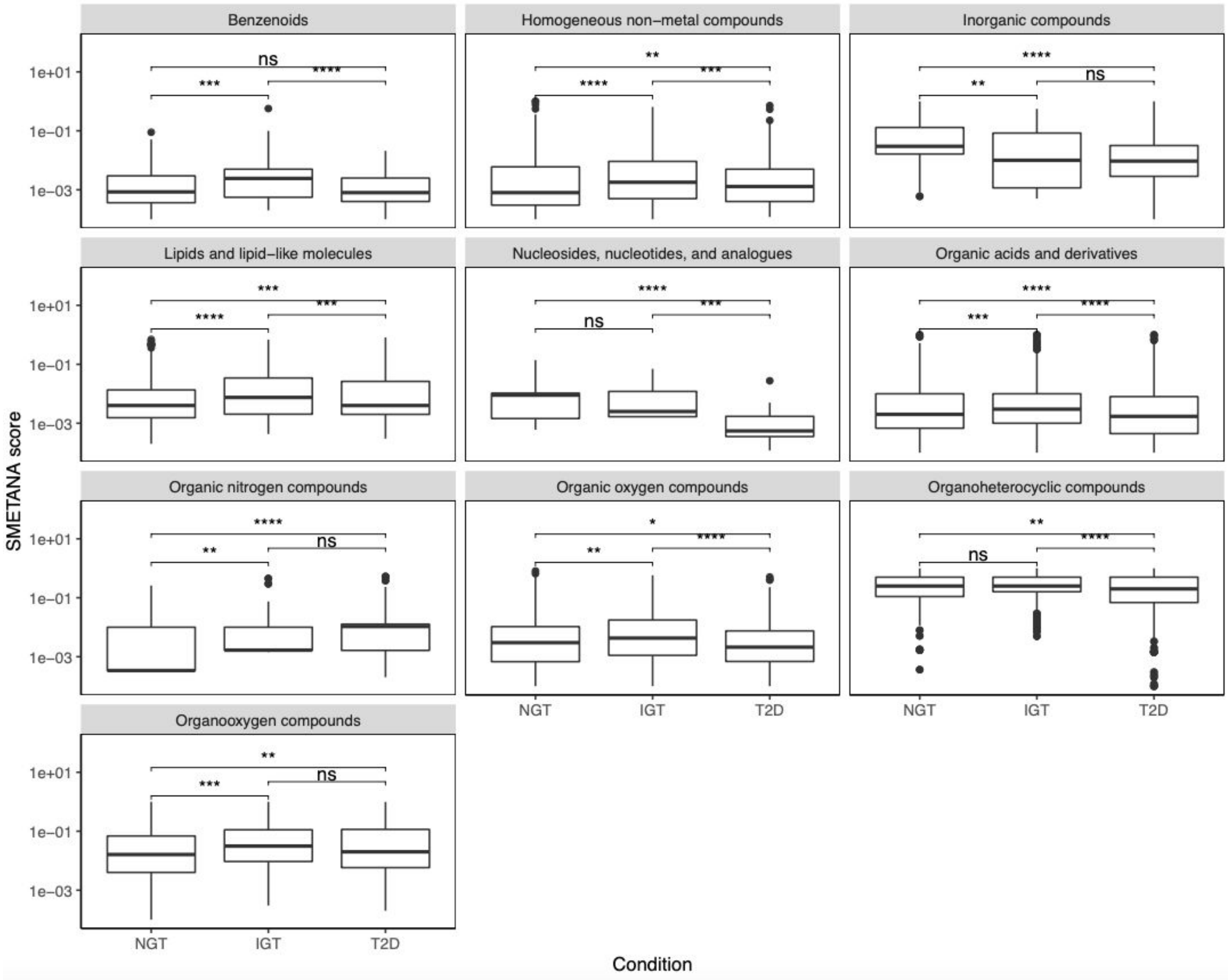


**Supplementary figure 12.** Distribution of SMETNA scores for the 22 metabolites with statistically significant differences across conditions grouped by broad metabolite class.

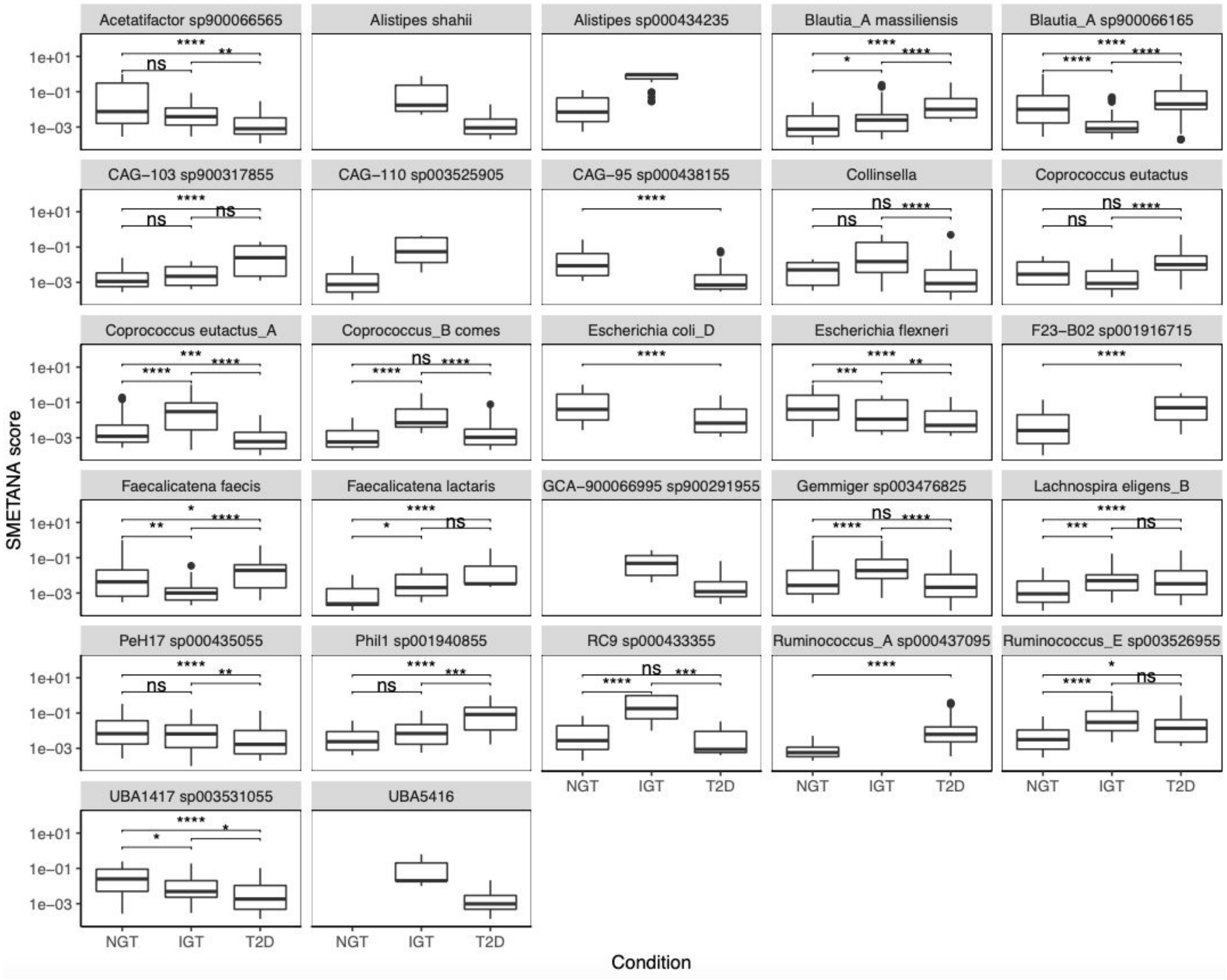


**Supplementary figure 13.** Distributions of SMETANA scores for 27 donor species where at least one comparison between disease groups yielded a highly statistically significant difference (p-value < 0.0001).

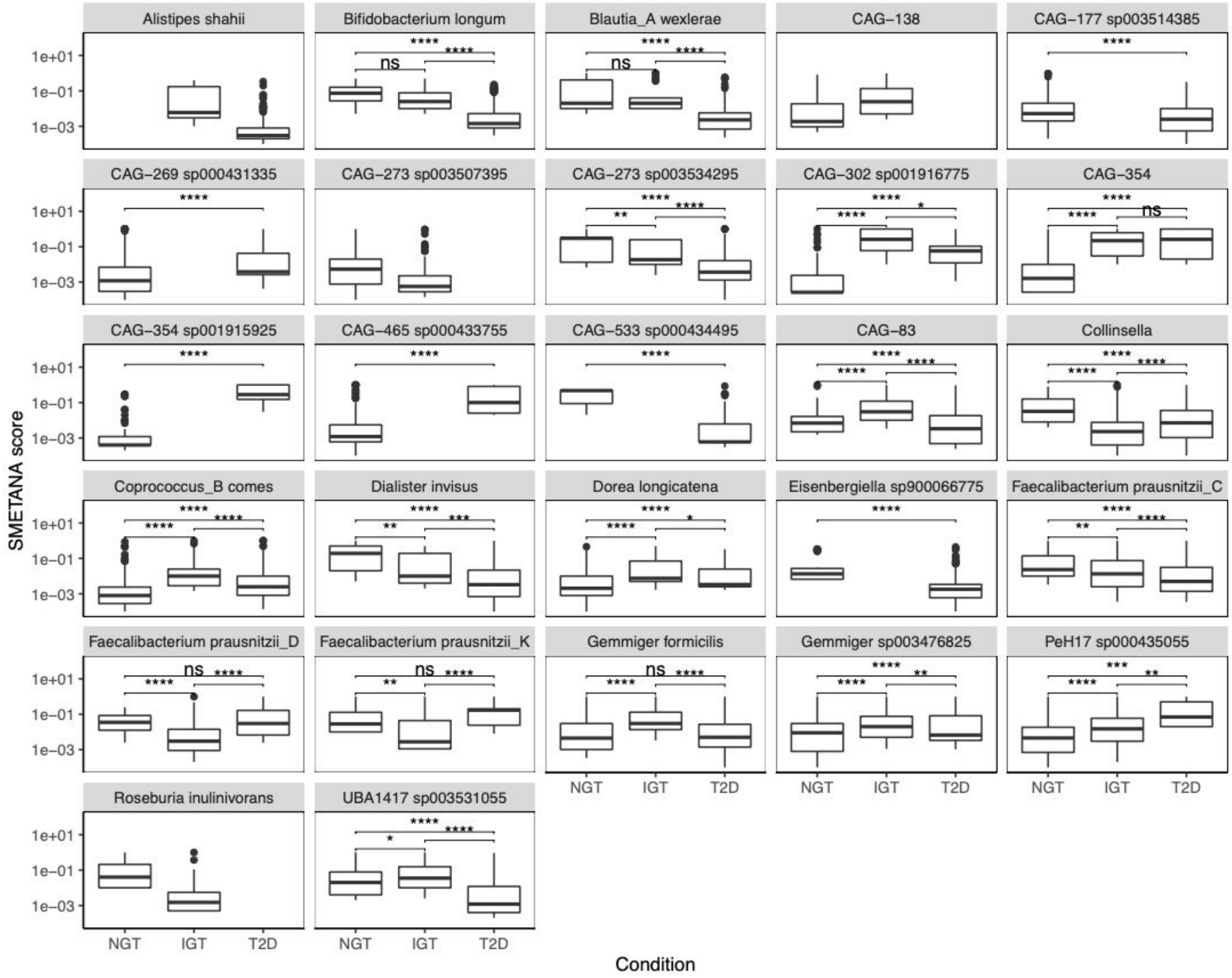


**Supplementary figure 14.** Distributions of SMETANA scores for 27 receiver species where at least one comparison between disease groups yielded a highly statistically significant difference (p-value < 0.0001)
